# *Rhizocarpon geographicum* lichen discloses a highly diversified microbiota carrying antibiotic resistance and Persistent Organic Pollutants tolerance

**DOI:** 10.1101/2022.07.25.501376

**Authors:** Alice Miral, Adam Kautsky, Susete Alves-Carvalho, Ludovic Cottret, Anne-Yvonne Guillerm-Erckelboudt, Manon Buguet, Isabelle Rouaud, Sylvain Tranchimand, Sophie Tomasi, Claudia Bartoli

**Author notes:** **To whom correspondence may be addressed to:** Claudia Bartoli. **Competing Interest Statement:** The authors declare no competing interests.

## Abstract

As rock-inhabitants, lichens are exposed to extreme and fluctuating abiotic conditions associated with poor sources of nutriments. These extreme conditions confer to lichens the unique ability to develop protective mechanisms. Consequently, lichen-associated microbes disclose highly versatile lifestyles and ecological plasticity, enabling them to withstand extreme environments. Because of their ability to grow on poor and extreme habitats, bacteria associated with lichens can tolerate a wide range of pollutants and on the other hand secrete antimicrobial compounds. In addition, lichen-associated bacteria were described to harbor ecological functions crucial for the evolution of the lichen holobiont. Nevertheless, the ecological features of lichen-associated microbes are still underestimated. To explore the untapped ecological diversity of lichen-associated bacteria, we adopted a novel culturomic approach on the crustose lichen *Rhizocarpon geographicum*. We sampled *R. geographicum* in French habitats exposed to oil spill and we combined 9 culturing methods with 16S rRNA sequencing to capture the largest bacterial diversity. A deep functional analysis of the lichen-associated bacterial collection showed the presence of a set of bacterial strains resistant to a wide range of antibiotics and displaying tolerance to Persistent Organic Pollutants (POP). Our study is a starting point to explore the ecological features of the lichen microbiota.

## Introduction

The term symbiosis was first introduced by Albert Bernhard Frank in 1876 to describe the mutualistic association between a fungal partner and a photobiont leading to lichen symbiotic organisms [1]. Lichens are widely distributed and have an incredible adaptability to highly contrasted ecosystems. They are pioneer species colonizing perturbated and extreme habitats and they display tolerance to dryness, UV-exposition, pollutants etc. [2]. Despite the extraordinary ecological features harbored by lichens, studies on their ecology and evolution are still in their infancy and mostly addressed on the fungal partner. Nevertheless, a recent work from Keller *et al*., 2022 [3] focuses on the impact of algae genomic diversity to explain the evolutionary trajectories of a wide range of terrestrial and aquatic lichens. A third partner, the lichen-associated-microbiota, not strictly associated with the symbiotic interaction, has been described as a key element of the lichen evolutionary history [2, 4–6]. By consequence, lichens have been reconsidered as holobionts [1, 7, 8]. The holobiont is a theoretical concept describing the host and its microbiome as unique evolutionary entity [9]. Looking at lichens as holobionts offers new perspectives on how to explore the lichen-associated biodiversity and their associated ecological functions [10].

Since the beginning of the 20^th^ century, lichen-associated bacteria were described to harbor ecological functions crucial for the maintenance of the lichen symbiotic system [5, 11–14]. Among the ecological functions, lichen-associated bacteria display resistance activity against compounds produced by the lichenic mycobiont. For instance, bacteria associated to lichens are known to resist to the usnic acid produced by several lichen species [15] and exhibiting a strong antimicrobial activity toward a wide-range of microbes [5, 16, 17]. In addition, bacteria isolated from seven lichen genera showed antibacterial activities against bacterial human pathogens [18]. Based on these previous studies, we can assume that lichen-associated bacteria are potential candidates in producing important and novel antibiotic compounds. Therefore, it is urgent to explore under a both ecological and biotechnological context, the diversity and the functional properties of microbial communities harbored by lichens.

In order to dissect the ecological functions behind the microbe-lichen interaction, the culturable and the unculturable microbiota needs to be deeply explored. Studies on the lichen microbiota are mostly based on molecular fingerprints [5, 19, 20], molecular cloning approaches [21] or 16S rRNA gene Illumina sequencing [22]. High-throughput sequencing methodologies (i.e. metabarcoding or metagenomics) are undoubtedly powerful to draw a whole picture of the lichenic microbiota. Conversely, they suffer from important limitations when applied to functional ecology [10]. Firstly, depending on the sequencing depth, microbial species can only be detected above a certain threshold. By consequence, low-abundance species with central ecological functions in the holobiont maintenance can be underestimated or undetected [23]. Secondly, functional studies aiming at understanding the ecological interaction regulating the lichen holobiont are dependent to deep microbial isolation efforts. Because many host-associated microbes are uncultivable outside their habitat of origin, improving cultivability is a prerequisite for a better understanding of metacommunities interactions and their functions in complex biotic systems [10, 24, 25]. Culturomics is a high-throughput culturing approach combining culture methods with mass spectroscopy or 16S ribosomal RNA sequencing in order to isolate and taxonomically affiliate a wide microbial diversity. Culturomics have been extensively employed in the human-gut microbiome field by expanding our knowledge on the bacterial repertoire colonizing the human gut [26]. Nevertheless, this promising approach has been unfortunately poorly utilized outside the medical field and even in a less extent on lichens [27].

Here we adopted a culturomic approach on *Rhizocarpon geographicum*, a crustose lichen pioneer of exposed rock surfaces and the first terrestrial substrate available for living organisms on Earth [28, 29]. We sampled *R. geographicum* in French zones strongly affected by hostile environmental conditions (wind, sea sprays, snow). Moreover, three of the sampling locations had history of oil spills. As a previous study revealed that the interaction of mutagenic compounds in natural environments is correlated to the development of antibiotic resistance genes in the bacteria [30], we investigated the possible link between antibiotic resistance and tolerance to Persistent Organic Pollutants (POP). We used 9 culturing media mimicking the lichen habitat and two culturing methods to isolate the widest bacterial diversity. We amplified a portion of 16S rRNA gene to affiliate the bacterial species composing *R. geographicum* microbiota. We pointed out the presence of a set of bacterial strains resistant to a wide range of antibiotics and displaying tolerance to two POP: the methyl *tert*-butyl ether (MTBE) and the perfluorooctanoic acid (PFAO). Our functional-ecological study on the *R. geographicum*-associated bacteria stresses the need to consider lichens as both source of important ecological functions and reservoirs of biotechnologically relevant bacterial species which can be used as markers of the habitat health.

## Material and methods

### *Collection of* Rhizocarpon geographicum *populations*

*R. geographicum* samples were collected in January 2021, under specific municipality authorizations, in 6 sites situated in France (Fig.1a) on: i) the Atlantic coastal area (Ille-et-Vilaine, Finistère and Côtes d’Armor - Brittany) ii) the English Channel coastal area in the northern limit of the Mont-Saint-Michel in Carolles (La Manche, Normandy), iii) the island area in Baulon and Plounéour-Ménez (Ille-et-Vilaine et Finistère) and iv) low mountains of Itxassou (Pyrénées Atlantiques, Nouvelle Aquitaine). Sampling localities were selected based on the habitat diversity (Fig. 1b, c and d). Three of these geographic sites (Finistère, Côtes d’Armor et La Manche) were chosen because they had a history of oil spills. For each location (called *R. geographicum* population), lichen samples were collected randomly on four rocks (Supplementary Table S1) and rock fragments were transported in sterile Petri dishes stored in individual plastic bags and processed within 6 hours post collection.

**Figure 1.**
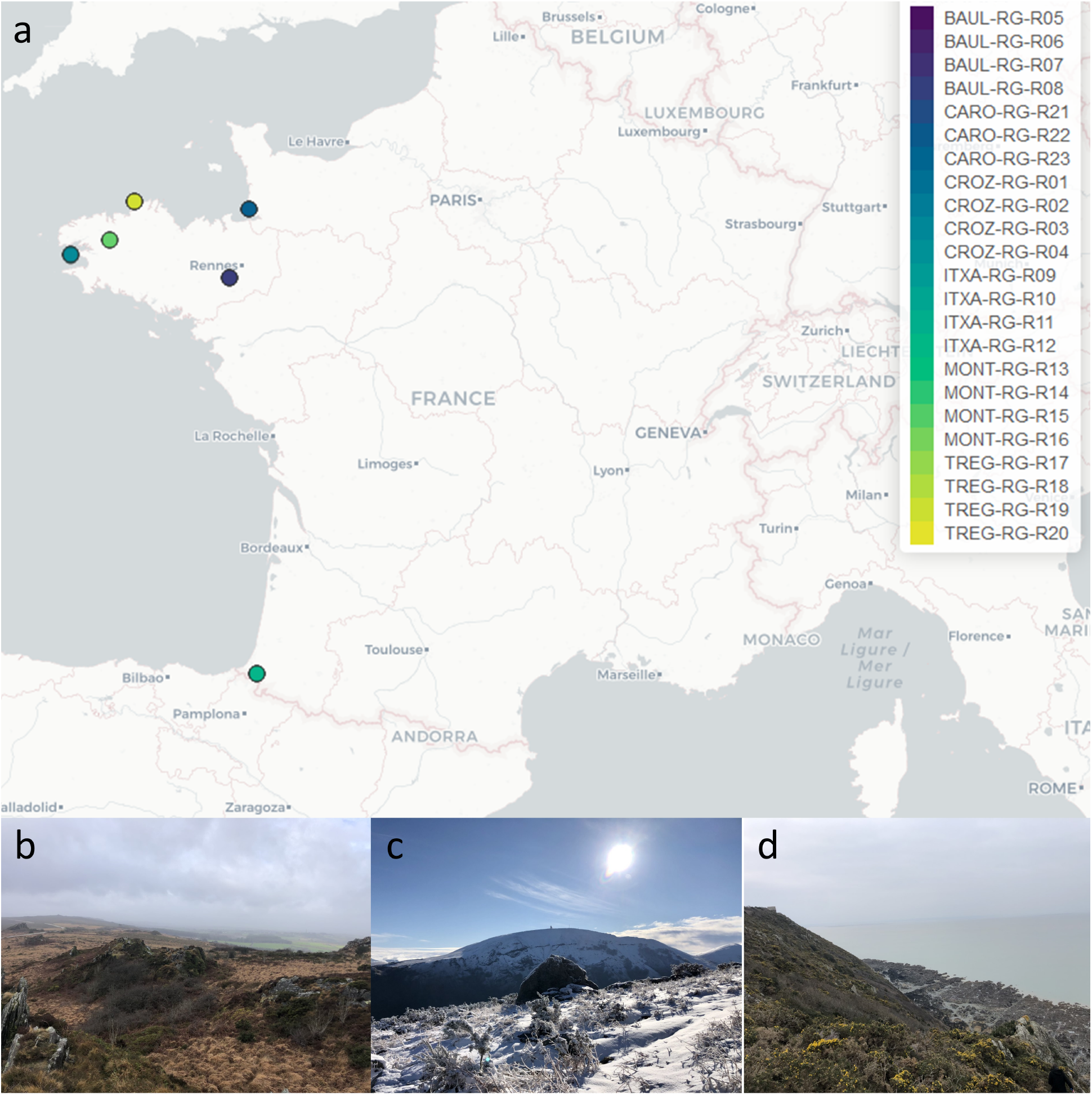
*Rhizocarpon geographicum* populations. (a) locations where populations were sampled across France (north and south). Colored dots represent the *R. geographicum* populations that were collected in the 6 sites, the color gradient within the same localities indicates the different harvest locations. Four populations per site were sampled. (b) representing MONT populations, (c) ITXA populations, and (d) CARO populations diversity of the terrestrial and maritime habitats harboring *R. geographicum*.

### Media used for culturomics

Three types of matrices were prepared as organic nutrient source and were supplemented into the media. For the first type of matrix, we developed a lichen filtrate. For this 50 mg of *R. geographicum* were scrapped and mixed with 100 mL of seawater collected from the site of sampling. The suspension was centrifuged for 10 min at 5000 rpm at 4°C. The supernatant was recovered and sterilized by filtration through 0.22 µm pore-size Millipore^®^ membranes. The second type of organic matrix was performed on a macroalgal filtrate. For this, 155 g of *Ascophyllum nodosum* and 95 g of *Laminaria digitata* collected at la Pointe de Crozon (Britany, France) were rinsed with seawater. These algae co-habit in the ecosystem with *R. geographicum* and are part of the lichen natural habitat. Macroalgae were roughly cut before being homogenized with a clean kitchen blender then bag-mixed with a Stomacher. From the concentrated solutions, 400 mL were recovered and filled with 600 mL of seawater collected at la Pointe de Crozon, previously sterilized by filtration. Once crushed, the algal pastes were passed through a stamen. Filtrates were then centrifuged for 10 min at 5000 rpm at ambient temperature. Filtrates were then filtered through 0.22 µm pore-size Millipore^®^ membranes to guarantee sterility. The third organic matrix consisted in a microalgal filtrate. *Coccomyxa viridis*, a microalgal associated with *R. geographicum*, collected on the Atlantic coastal area in Crozon, was previously isolated on the International Streptomyces Project 2 medium (ISP2, dextrose 4 g.L^-1^ [Sigma-Aldrich, St. Louis, MO, USA], yeast extract 4 g.L^-1^ [Sigma-Aldrich, St. Louis, MO, USA], malt extract 10 g.L^-1^ [Sigma-Aldrich, St. Louis, MO, USA], agar 15 g.L^-1^ [Sigma-Aldrich, St. Louis, MO, USA]). The total mass present in one Petri dish of *C. viridis*, was added into 50 mL of NaCl 0.85 % then transferred into 500 mL of ISP2 liquid medium. Culture was maintained under agitation (120 rpm) for 2 weeks. After this growing time, 500 mL of sea water was added into the *C. viridis* suspension and sterilized by filtration as described above.

The three lichen, macroalgal and microalgal-based matrices were then used to develop culturing media (Supplementary Methods). The media codes indicated in the Supplementary Methods are part of a largest list of culturing media developed at the IGEPP Laboratory (INRAE, France) with the aim to optimize culturomics on several substrates. Medium 17B (lichen microalgal-based minimal medium) was used for bacterial enrichment and a total of 9 media were used for bacterial isolation (Supplementary Methods). Three media (18F, 19F and 20F) based on lichen and/or algal suspensions were supplemented by 100 mg.L^-1^ of chloramphenicol, 300 mg.L^-1^ of streptomycin and 100 mg.L^-1^ of penicillin G to isolate antibiotic resistant strains. Detailed protocols for media preparation are listed in the Supplementary Methods.

### Bacterial isolation, characterization and taxonomic affiliation

The 24 crustose lichen samples were aseptically scrapped using a sterile scalpel and placed into sterile 15 mL Falcon^®^ tubes containing 5 mL of sterilized distilled water. The fresh mass of each lichen used for isolation is listed in Supplementary Table S2. To isolate the larger bacterial diversity, we adopted two methodologies. Firstly, dilutions (from 10^−1^ to 10^−3)^ were directly plated on the 9 media (media supplemented with 100 mg.L^-1^ of cycloheximide: 01B, 08B, 14B, 15B; 16B and 17B and media supplemented with 100 mg.L^-1^ of chloramphenicol, 300 mg L^-1^ of streptomycin and 100 mg.L^-1^ penicillin G18F, 19F and 20F) described in the Supplementary Material. The samples were incubated under dark conditions at 15 and 20°C until colonies appeared for 6 weeks. Secondly, the 24 lichen samples were enriched into 500 mL Erlenmeyer containing 50 mL of microalgae filtrate and 500 µL of lichen filtrate. The four samples for each *R. geographicum* population were pooled and 100 µL of each lichen suspension were added to the enrichment broth. Erlenmeyer flasks were sealed with a cotton and aluminum cap then maintained under agitation (100 rpm) at 15°C for 4 weeks. Once a week, the cap was aseptically removed for 15 minutes to aerate the cultures. After 4 weeks, dilutions from the enrichment solutions were plated in triplicate on the 9 media (Supplementary Material). Plates were incubated under dark conditions at 20°C for 6 weeks.

For both experimental procedures (direct plating and enrichment), colonies showing distinct morphologies were purified on the medium of origin and stored at -20°C into 30 % of glycerol. Pure colonies were placed into 96-well plates containing a DNA stabilizing buffer composed by 10 mM TRIS, pH 8.0, 0.1M EDTA, pH 8.0 and 0.5 % SDS. Plates were stored at -20°C prior DNA extraction that was performed with the protocol described in Vingataramin and Frost (2015) [31], with minor modifications. Briefly, 300 µL of EtNa DNA extraction reagent were added to preheated DNA plates containing the stabilizing buffer. The mix was heated at 90°C for 10 min and centrifuged at 4000 rpm for 30 min. The supernatant was removed and the pellet resuspended in 100 µL of DNA suspension solution. Bacterial identification was performed by amplifying a region of the 16S as described in the Supplementary Material.

### Characterization of antibiotic resistant strains

Twenty-four bacterial strains (Supplementary Table S3) able to grow on media containing antibiotics (Supplementary Methods), were tested for their antibiotic resistance ability against 12 antibiotics (Supplementary Table 4). For this, the 24 strains were incubated in 10 mL of Tryptic Soy Broth (TSB) (Sigma, Reference T8907-1KG) from 2 to 4 days at 20°C under agitation (120 rpm). Strains were then tested at both exponential (*exp*) and stationary (*sta*) phase. After incubation, 10 µL of each strain at *exp* or *sta* phase were inoculated into sterile 96-well plates containing 90 µL TSB and the appropriate antibiotic concentration (Supplementary Table S4). TSB in absence of the antibiotic was used as control. Three independent experiments (temporal blocks) were performed and for each experiment three replicates were included. Plates were incubated for 72 h at 20°C under agitation (120 rpm) then the optical densities (OD) were measured at λ = 620 nm with a microplate reader Multiskan™ FC, Thermo Scientific™. The 9 values for each *strain* × *antibiotic* obtained from the 3 independent experiments were used to estimate the antibiotic effect on the bacterial growth on both *exp* and *sta* conditions. Also, *exp* and *sta* conditions were nested into the analysis to estimate their effect on strain growth variability. For this, we first estimated an antibiotic coefficient by dividing the OD of each strain growing in presence of the antibiotic by the OD of the strain growing in absence of the antibiotic. Coefficient values were integrated into a generalized-linear-mixed model that was run by using the glmer function implemented in lsmeans and lme4 R packages [32, 33]. Replicates and the temporal blocks (i.e. the 3 independent experiments) were integrated in the model as random effects. *P*-values were corrected for FDR and barplots were built using the ggplot2 and reshape2 packages [34, 35] on the lsmeans.

Nine strains showing a high antibiotic resistance range were whole-genome sequenced (Supplementary Table S3) and analyzed as described in the Supplementary Methods. Two strains that were not identified by 16S sequencing (CARO-RG-8B-R23-01 and CARO-RG-8B-R24-01) were also whole-genome sequenced to investigate their taxonomical affiliation.

### Tolerance to persistent organic pollutants

A sub-set of 394 bacterial strains belonging to unique 16S clusters and showing a suitable growth rate when re-cultured on TSA medium after storage, were tested for their ability to tolerate and grow in presence of Persistent Organic Pollutants (POP). For this, the 394 strains where grown for 3-days on TSA and then inoculated into TSB supplemented either with perfluorooctanoic acid (PFOA) at a final concentration of 20 mg L^-1^ or methyl *tert*-butyl ether (MTBE) at a final concentration of 2 g L^-1^. Controls consisted of strains inoculated on TSB only. POP screening was performed in triplicates on 96-well plates containing a volume of 200 µL of TSB supplemented with the POP. Each well was inoculated with one pure bacterial colony. The inoculated 96-well plates were incubated for 7 days at room temperature then the OD were measured at λ = 620 nm with microplate reader Multiskan™ FC, Thermo Scientific™. Experiment was independently repeated two times (temporal blocks) and each block was constituted by three strain replicates. The 9 values for each *strain* obtained from the 3 temporal blocks were used to estimate the POP tolerance. For this, we estimated a POP tolerance coefficient by dividing the OD of each strain growing in presence of the given POP by the OD of the strain growing in absence of the POP. Coefficient values were integrated into a generalized-linear-mixed model that was run by using the *lmer* function implemented in lsmeans and lme4 R packages [32, 33]. The 9 replicates were considered in the model as random effects. A threshold of POP tolerance coefficient >1 was considered as in index of bacterial growth on the POP.

## Results

### *Culturomics revealed a* Rhizocarpon geographicum *highly diversified microbiota*

*R. geographicum* was sampled in 6 highly diversified maritime and terrestrial habitats located in France (Fig. 1a, 1b). These habitats were selected to maximize niche diversity but also to include French littorals with a history of oil spill [36]. For this, Crozon and Trégastel located in Brittany, and Carolles located in Normandy were chosen to collect *R. geographicum* associated microbes potentially harboring tolerance to hydrocarbons and pollutants. To capture the highest bacterial diversity, we applied a culturomics approach by using 9 isolation media (Supplementary Methods) and two culturing methods: direct plating and enrichment with microalgae prior plating. In order to increase the recovery of the bacterial communities, we mimicked the nutritional and environmental conditions of *R. geographicum*. Seven of the selected media consisted in algae-based and/or lichen-based media and they were developed in our study to reproduce the *R. geographicum* habitat and isolate recalcitrant bacterial species. Three media were supplemented with antibiotics to isolate bacterial strains displaying resistance to a wide range of antibiotics. We attempted to isolate antibiotic resistant bacteria, as Grube and collaborators identified in lichen-associated bacteria genes coding for multidrug resistance efflux pumps [11, 37]. Moreover, we used antibiotics because there are molecular evidences that the presence of POP can be implicated in the selection of microbes displaying antibiotic resistance genes in contaminated soils [30].

We isolated a total number of 1,913 bacteria from *R. geographicum* samples. Among the 1,913 we succeeded to amplify the 16S rRNA of 1, 063 strains. Based on the 16S sequences, 364 bacterial clusters showing 100% of sequence homology with at least one strain were identified thought the CD-HIT software [38] (Supplementary Data Set 1 and Data Set 2). The remaining 699 unique bacterial clusters (i.e. bacterial strains showing a unique 16S sequence) where taxonomically affiliated (Supplementary Data Set 2). Bacterial abundance (i.e. number of bacterial isolated per each medium divided by the total number of bacteria) showed a heterogenous success to isolate bacteria across the isolation media used (Fig. 2). The highest number of strains were recovered on the 01B (Nutrient Agar) and the 08B (TSA) media (Fig. 2). A lower number of bacterial strains were recovered from the algae or lichen-based media (14B, 15B, 16B, 17B) and few antibiotic resistance strains (N = 126) were isolated on the 18F, 19F and 20F media (Fig. 2). Similar results were obtained when analyzing the abundance of distinct bacterial isolated species on each medium (Supplementary Fig. S1). On the other hand, media designed to reproduce the lichen environment (14B, 15B, 16B, 17B, 18F, 19F and 20F, Supplementary Methods) allowed to isolate 17 bacterial species that were not recovered on the Nutrient Agar (01B) and TSA (08B) medium and that where specific to the isolating medium (Table 1). Most of the species isolated from media reproducing the lichen environment are known to evolve in water reservoirs and polluted environments. For instance, *Aquaspirillum arcticum, Erythrobacter* sp, *Klenkia taihuensis, Paracoccus chinensis, Microterricola gilva*, and *Kocuria polaris* (Table 1), were previously isolated from marine environments [39–42], sediments [43, 44], seaweed [45] or cyanobacteria [46]. It is to note that various *Erythrobacte*r, *Paracoccus, Kocuria, Pseudomonas* or *Gordonia* species were already isolated from marine or maritime Brittany lichens and *P. helmanticensis* from an island Austrian lichen [42]. *Geobacillus* sp. and *Gordonia* sp. are described to grow on thermal area and polluted and contaminated environments [47–53]. *Geobacillus* sp. is also described to produce antibiotics [54]. Our results suggest that media reproducing the habitat of origin help in isolating rare species that can play important roles on the ecological network of the habitat.

**Figure 2.**
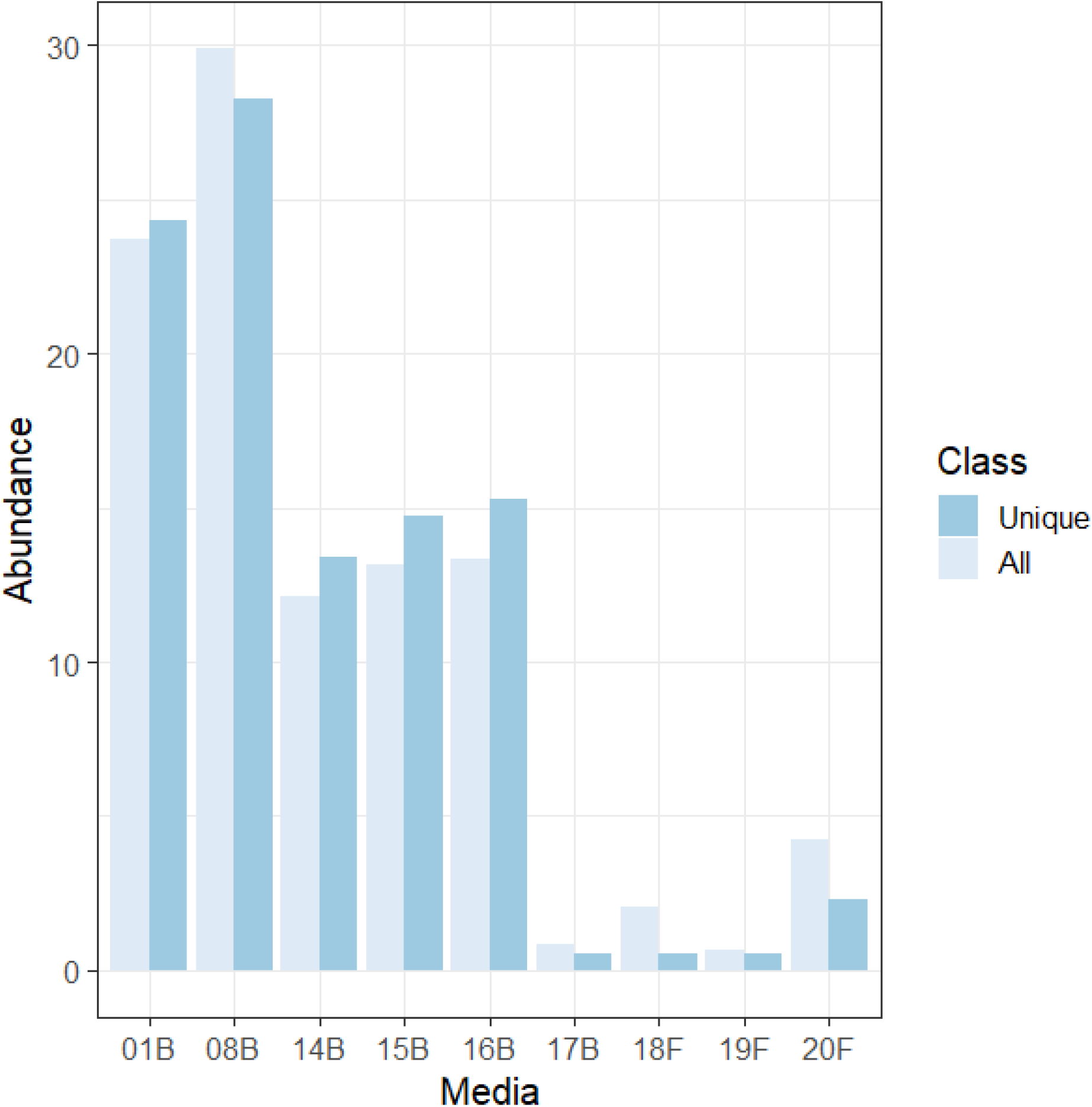
Number of bacteria strains isolated from each medium. Bar plot is based on the abundance of each bacterial strain (expressed in %), *y*-axis calculated by dividing the number of strains recovered on each medium by the number of the total strains. *x*-axis represent the isolation medium used. The class color scale indicates the overall isolated bacteria (light blue) and the unique bacterial clusters (dark blue) based on their 16S sequences after CD-Hist analysis.

**Table 1.** Bacterial species isolated specifically from lichen and/or algal-based media. Full media description is provided in Supplementary Methods.

| Bacterial species | Isolation medium | Medium description |
| --- | --- | --- |
| *Aquaspirillum arcticum* | 15B | Lichen algal-based medium |
| *Geobacillus* sp. | 16B | Lichen-algal based minimal medium |
| *Jeongeupia chitinilytica* | 15B | Lichen algal-based medium |
| *Klenkia taihuensis* | 14B | Macroalgae-based medium |
| *Kocuria polaris* | 14B | Macroalgae-based medium |
| *Microterricola gilva* | 16B | Lichen-algal based minimal medium |
| *Paracoccus chinensis* | 19F, 20F | Lichen-algal antibiotic based |
| *Pseudomonas gessardii* | 20F | Lichen-algal antibiotic based |
| *Pseudomonas helmanticensis* | 16B, 20F | Lichen-algal based medium & Lichen-algal antibiotic based |
| *Roseomonas mucosa* | 15B | Lichen algal-based medium |
| *Sinomonas* sp. | 15B | Lichen algal-based medium |
| *Skermanella* sp. | 14B | Macroalgae-based medium |
| *Tardiphaga* sp. | 14B | Macroalgae-based medium |
| uncultured *Erythrobacter* sp. | 16B | Lichen-algal based minimal medium |
| uncultured *Gordonia* sp. | 16B | Lichen-algal based minimal medium |
| uncultured *Nakamurella* sp. | 8B, 16B | TSA and Lichen-algal based minimal medium |
| *Zafaria cholistanensis* | 14B | Macroalgae-based medium |

Based on 16S taxonomy affiliation, the most abundant bacterial class isolated by culturomics belong to α-Proteobacteria followed by Actinomycetia, β-Proteobacteria and γ-Proteobacteria (Fig. 3a). Among the three most representative bacterial class, the α-Proteobacteria are mostly constituted by the Hyphomicrobiales and Sphingomonadales families (Fig. 3b). The Actinomycetia are mostly represented by the Micrococcales and the Frankiales orders (Fig. 3c). The β-proteobacteria are almost exclusively represented by the Burkholderiales order. These results are coherent with those obtained previously by a metagenomic study on different crustose lichens, including *R. geographicum* [20].

**Figure 3.**
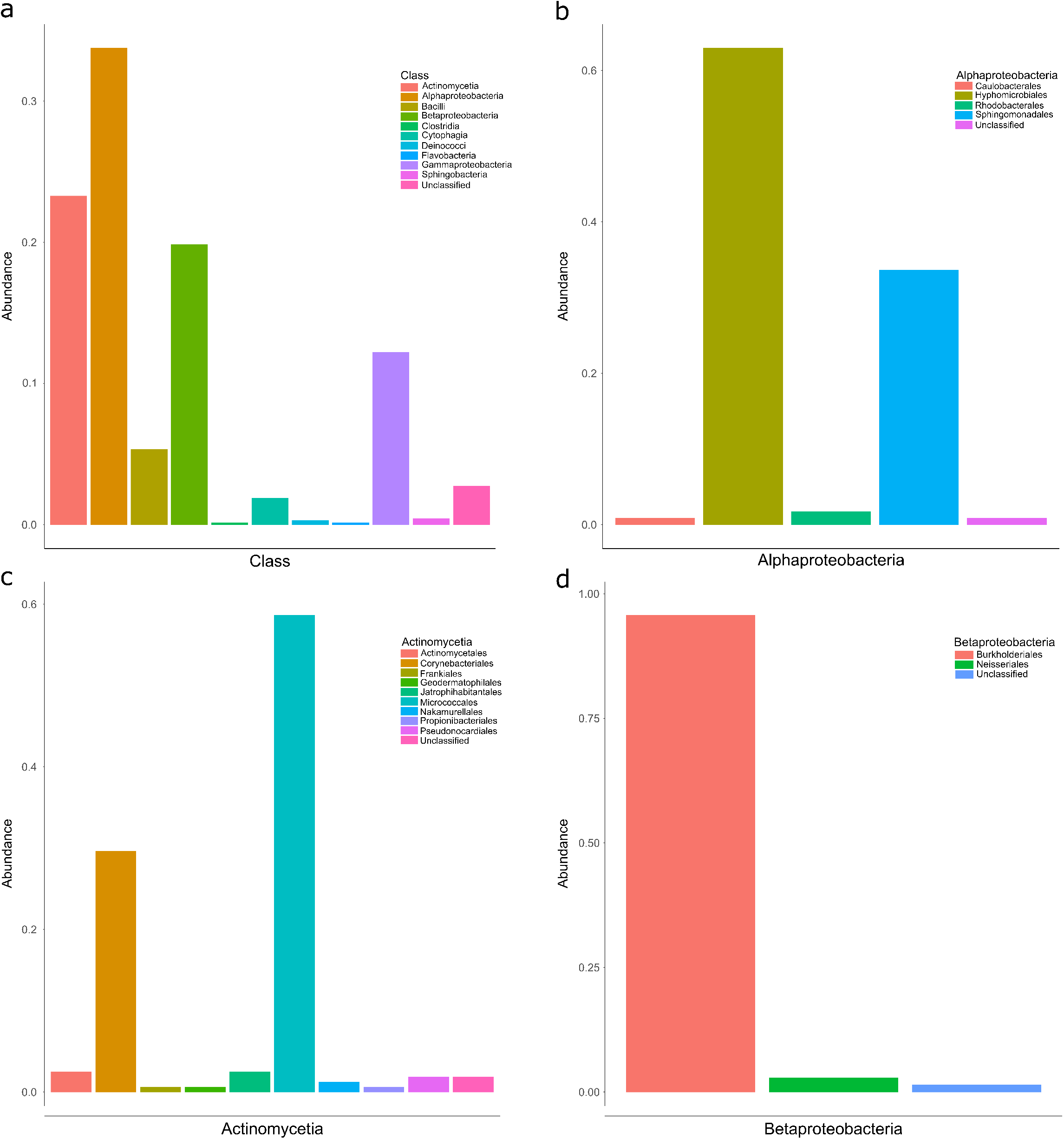
Abundance of bacterial class and families found on all media. (a) bar plot on the frequency (N° of each bacterial class/N° of total isolated strains) of the bacterial class found on *Rhizocarpon geographicum*. (b) bar plot on the frequency of the bacterial families included in the α-Proteobacteria class, (c) bar plot on the frequency of the bacterial families included in the Actinomycetia class and (d) bar plot on the frequency of the bacterial families included in the β-Proteobacteria class.

Three strains CARO-RG-8B-R24-01, CARO-RG-8B-R23-01 and MONT-RG-14B-R14-06 did not match with any 16S sequences in GenBank. We then extracted the total DNA of these isolates for whole-genome sequencing. For MONT-RG-14B-R14-06, DNA did not pass the quality test and was not sequenced. Statistics on the genome quality for the two genomes is reported in Supplementary Table S5. Phylogeny on the whole genome performed by the Type Strain Genome Server (TYGS) https://tygs.dsmz.de/ showed that CARO-RG-8B-R23-01 strain belongs to the *Microbacterium lacticum* species, on the other hand CARO-RG-8B-R24-01 belongs to a potential new species of the *Sphingomonas* genus (Supplementary Fig. S2) distantly related to *Sphingomonas glacialis* a psychrophilic Gram-positive bacterium isolated previously on alpine cryoconite [55].

### Culturomics allows to isolate taxonomically diversified antibiotic resistant bacterial species

We identified 126 bacterial strains from media containing antibiotics. Among the 126, only 87 were selected for their 16S sequence quality (Supplementary Data Set 3). Based on the phylogenetic diversity obtained on a fragment of the 16S sequence (Supplementary Fig. S3) we selected 24 strains (Supplementary Table S3) for testing their antibiotic resistance degree against 12 antibiotics (Supplementary Table S4) used as therapeutics. Strains were selected to maximize their phylogenetic diversity and geographic origin. Among the 24 bacterial isolates, 10 belonged to the *Sphingomonas* genus, 8 to the *Pseudomonas* genus, 3 strains were affiliated to the *Salinarimonas* genus, one strain was identified as *Paracoccus chinensis*, one was affiliated to the actinomycete *Amycolatopsis panacis* and one strain belonged to the *Janthinobacterium* genus. Linear-mixed model performed on the antibiotic resistance coefficient showed a high effect of the Strain *factor* for the antibiotic response (Supplementary Table S6), by confirming the heterogenicity of strains in response to the tested antibiotics (Fig. 4). The *exp* and *sta* determining the Phase *factor* (whether strains have been inoculated at the exponential – *exp* – or stationary – *sta* – phase) was significant on 5 of the 12 antibiotics tested (gentamicin, kanamycin, vancomycin, colistin and cefalexin, Supplementary Table S6). On the other hand, a nested effect Strain × Phase *factor* was found for all antibiotics except penicillin G (Supplementary Table S6). This result confirms that the bacterial phase at the time of strain inoculation can have an impact on the strain resistance on several antibiotic classes (Fig. 4). Globally, as showed in Fig. 4, resistance to antibiotics was stronger when strains were inoculated at the stationary phase. Few strains were resistant to gentamicin, kanamycin and streptomycin (that are aminoglycoside class antibiotics inhibiting the 30S bacterial ribosome subunit synthesis). For vancomycin (a glycopeptide antibiotic inhibiting the bacterial cell wall biosynthesis), strains showed resistance only when inoculated at the stationary phase (Fig. 4).

**Figure 4.**
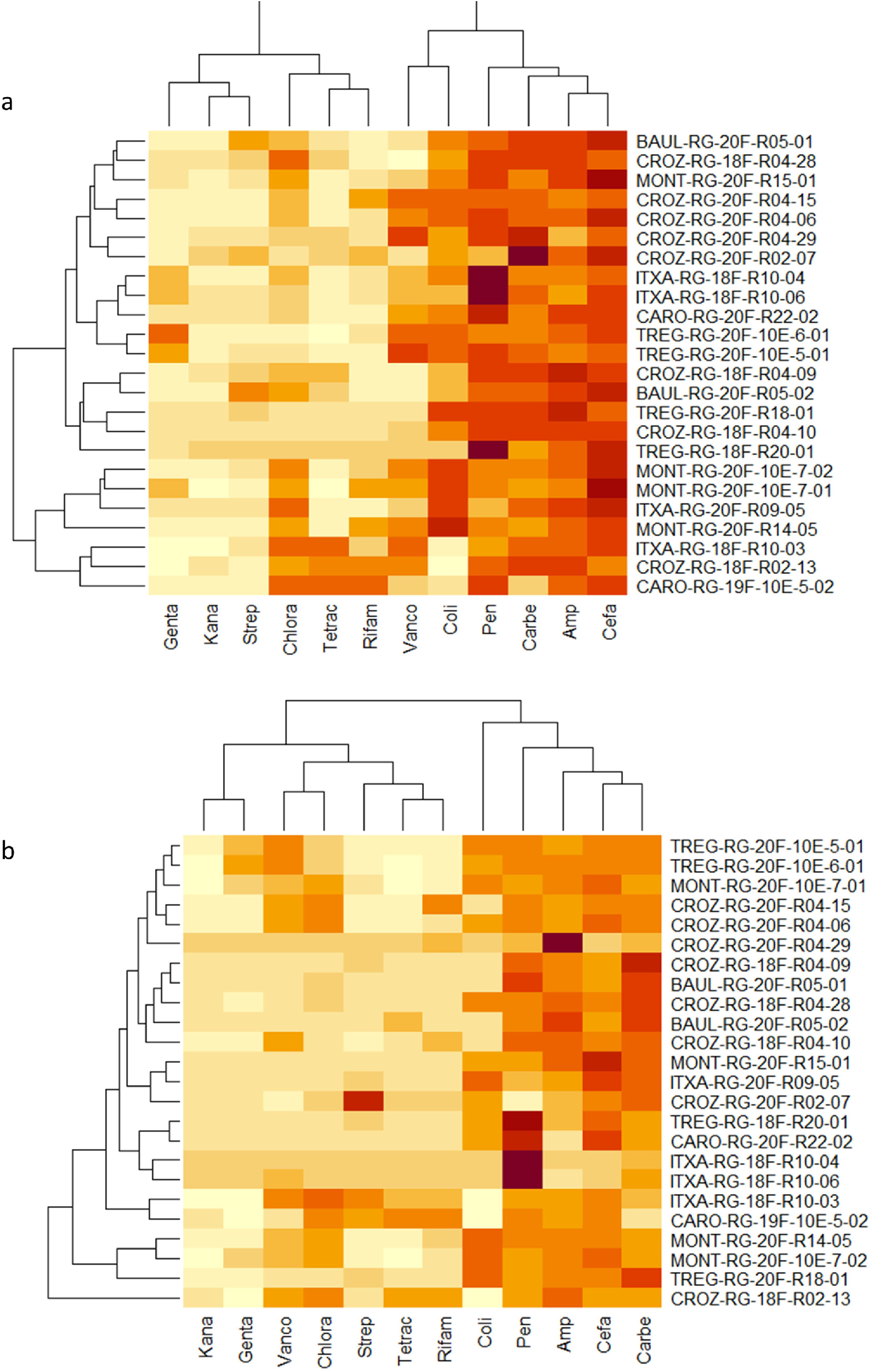
Heatmaps on the antibiotic resistance coefficient. Heatmaps were performed on the lsmeans obtained by running a liner-mixed model on the antibiotic resistance coefficient and by controlling for the block and replication effect. (a) heatmap on the antibiotic coefficients obtained when strains where inoculated at the stationary phase; (b) heatmap on the antibiotic coefficients obtained when strains where inoculated at the exponential phase. Name of the tested strains are reported in the *y*-axis, antibiotics are reported in the *x*-axis. Abbreviations: Genta = gentamicin, Kana = kanamycin, Vanco = vancomycin, Chlora = chloramphenicol, Tetrac = tetracycline, Rifam = rifampicin, Strep = streptomycin, Coli = colistin, Pen =penicillin G, Carbe = carbenicillin, Amp = ampicillin, Cefa = cefalexin. Color scale from dark red to light yellow represents high antibiotic resistance (dark red) and no resistance (light yellow).

### Whole-genome sequencing of antibiotic resistant bacteria suggests potential novel species harbouring Antimicrobial Resistance (AMR) genes

We sequenced and analysed 9 bacterial strains showing the highest degree of resistance to the tested antibiotics. Phylogenetic affiliation of the strains performed on the TYGS database [56], suggested that the 9 strains are potential novel species. The TYGS database is updated daily, however, type strains closely related to the sequenced strains could be missing. Based on the phylogenetic inference on the whole-genomes (Fig. 5a, 5b), the 9 potential new species are distantly related to: i) *Pseudomonas crudilactis* for the CROZ-RG-20F-R04-06 and CROZ-RG-20F-R04**-**15 strains ii) *Pseudomonas antartica* for TREG-RG-20F-10-E-5-01, TREG-RG-20F-10-E-6-0, MONT-RG-20F-10-E-7-02 and MONT-RG-20F-R14-5 strains iii) *Sphingomonas glacialis, Sphingomonas panacis* and *Sphingomonas ginsenosidivorax* for TREG-RG-20F-R18-01, BAUL-RG-20F-R05-02 and CROZ-RG-20F-R02-07 strains respectively (Fig. 5a, 5b). The closely related strains were isolated from various habitats. Indeed, *P. antarctica* and *S. glacialis* are psychrophile bacteria isolated from Antarctica water bodies [57] and Alpine glacier cryoconite respectively [55]. Both *P. antartica* and *S. glacialis* inhabit extreme environments and were described to carry antimicrobial activity [55, 58]. *P. crudilactis*, firstly isolated from raw milk, was shown to be resistant to several antibiotics [59]. *S. ginsenosidivorax*, isolated from ginseng [60] did not exhibit antibiotic resistance in the previous studies but is able to bio-transform ginsenosides, possessing antimicrobial activities [61].

**Figure 5.**
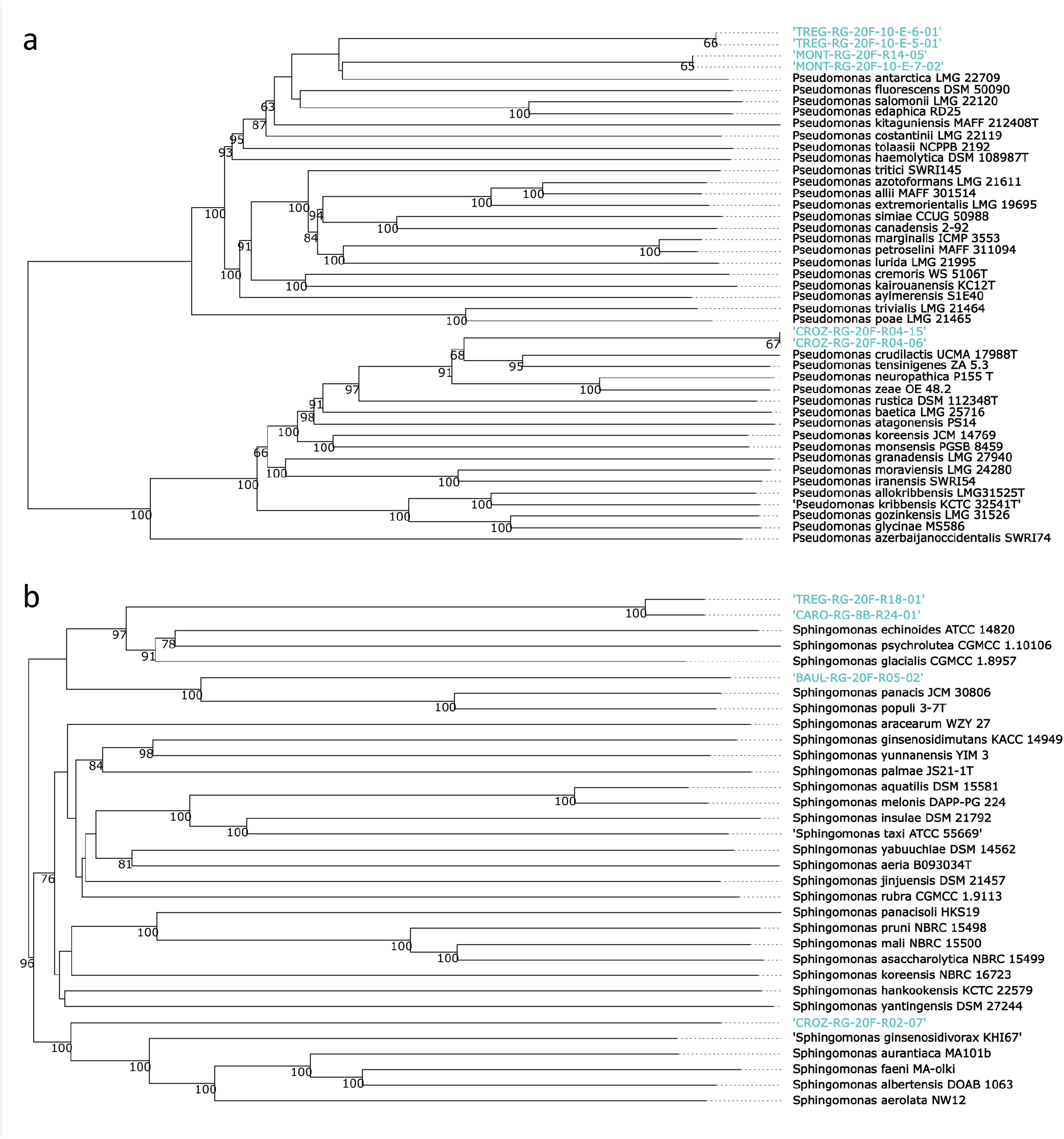
Phylogeny representing CROZ-RG-20F-R04-06, CROZ-RG-20F-R04-15, MONT-RG-20F-10-E-7-02, MONT-RG-20F-R14-05, TREG-RG-20F-10-E-5-01 and TREG-RG-20F-10-E-6-01 bacterial strains (a) and BAUL-RG-20F-R05-02, CARO-RG-8B-R24-01, CROZ-RG-20F-R02-07 and TREG-RG-20F-R18-01 (b) bacterial strains. Phylogenetic tree was inferred with FastMe 2.1.6.1 [88] on the TYGS platform [56] from GBDP distances calculated from whole-genomes. The tree was rooted ad midpoint. Tree was edited by iTOL [89]. Strains isolated from *R. geographicum* and the related tree branches are indicated in blue.

Analysis of the predicted KEGG metabolic pathways performed on the 9 genomes and the two genomes of the strains not affiliated by 16S, showed variability among the strains (Supplementary Data Set 4). The main variabilities reside in carbon degradation, nitrogen, sulfur and hydrogen cycles, vitamins and transporters as well as secretion systems. Cluster analysis based on the number of biological pathways and reactions showed similar results. On both number and type of pathways and reactions, all strains were clustered in agreement with their taxonomic affiliation (Fig. S4). The CARO-RG-8B-23-01 strains displayed a highly different metabolic profile that was quietly expected as belonging to an anaerobic bacterium (*Microbacterium lacticum*) (Supplementary Data Set 4, Fig. S4). Interestingly, the TREG-RG-20F-10E-6-01, TREG-RG-20F-10E-5-01, MONT-RG-20F-R14-05 and MONT-RG-20F-10-E-7-02 closely related to *P. antartica* (Fig. 5a) and carrying the strongest antibiotic resistance profile (Fig. 4) showed a similar metabolic profile (Figure S4, Supplementary Data Set 4)

To better understand the genetic bases of the antibiotic resistance activity of the 9 strains, we investigated the presence of Antimicrobial Resistance (AMR) proteins by using AMRFinder Plus [62] enabling accurate assessment of AMR gene content. We found that bacterial strains isolated from the Baulon lichen population carried class A β-lactamase and subclass B3 metallo-β-lactamase proteins (Supplementary Data Set 5). Strains from the Crozon lichen population displayed a wider range of AMR proteins that were annotated as efflux RND transporter permease subunit EmhB, efflux RND transporter outer membrane subunit EmhC, class C β-lactamase and copper resistance metal-translocating P1-type ATPase CueA (Supplementary Data Set 5). Strains from the Plounéour-Ménez population were found to carry NAD(+)-rifampicin ADP-ribosyltransferase, class C β-lactamase and fosfomycin resistance glutathione transferase proteins. Finally strains coming from the Trégastel population and showing the highest level of antibiotic resistance were found to harbour AMR proteins annotated as fosfomycin resistance glutathione transferase, class C β-lactamase, arsinothricin resistance *N*-acetyltransferase ArsN1 family B and APH(3’) family aminoglycoside *O*-phosphotransferase. Interestingly, the AMR profile was defined by the lichen population, suggesting that specific antimicrobial strategies are adopted at local/habitat level.

### Lichen bacterial strains harbour tolerance to persistent organic pollutants that is not correlated with the antimicrobial activity

In order to investigate the putative tolerance to persistent organic pollutants (POP) of strains of our collection and the possible connection with antibiotic resistance activity, perfluorooctanoic acid (PFOA) - a perfluoroalkyl substance found in aqueous film forming foams - and methyl *tert*-butyl ether (MTBE) - a petrol additive - were used as source of POP during bacterial growth on 394 strains (Supplementary Data Set 6). The 394 strains were selected because they were able to efficiently grow on TSA medium. Linear models demonstrated a highly significant effect of the *strain* factor (*χ*^2^ = 483.04, *P* = 0.001425) on MTBE tolerance and on the PFOA tolerant (*χ*^2^ = 590.87, *P* = 4.613e-10) by suggesting a high variability among the strains in the response to both POP tested (Supplementary Data Set 6). Regression on the LSmeans between the two POP tolerance coefficients showed a *R*^*2*^ = 0.11 suggesting a faint degree of correlation between both POP tolerance (Figure S5).

For the MTBE, within the 394 strains, 237 were able to tolerate the POP (tolerance coefficient ≥ 1) and 42 strains has a tolerance coefficient ≥ 1.5. The fifteen strains (3.8 % of the tested bacteria) showing the highest level of tolerance (coefficient ≥ 2) where recovered from the CARO (Carolles), CROZ (Crozon) and TREG (Trégastel) populations (Supplementary Data Set 6, Table 2), localities carrying a history of oil spill. This result strongly suggests that for the MTBE the ability of the strains to resist to the POP is linked to the habitat of the origin (Table 2). The 15 most tolerating strains belonged to 7 bacterial genera/species: i) *Frondihabitans* an actinobacterium known to colonize lichens ii) *Arthrobacter* an ubiquitous bacterial species colonizing several substrates iii) *Paenibacillus* a lichen-associated bacterium, iv) *Amantichitinum* a soil bacterium, v) *Deinococcus alpinitundrae* isolates previously from alpine environments and known to be resistant to ionizing radiation and vi) five strains belonged to the *Pseudomonas* and *Sphingomonas* genus. The species affiliation of the highest tolerant strains indicates that most of this strain are adapted to the lichen environment and are known to resist to extreme environments.

**Table 2.** Total number of tested strains on POP and % of strains tolerant to both MTBE and PFOA for each lichen population when considering a tolerant coefficient ≥ 2. Na indicates that any strain from the specific locality was identified with a tolerant coefficient ≥ 2. Localities indicated in bold correspond to habitats that have been subjected to oil spills in their history

| **Lichen population** | **Locality** | **% of tolerant strains for MTBE** | **% of tolerant strains for PFOA** |
| --- | --- | --- | --- |
| BAUL | Baulon | na | 0.76 |
| **CARO** | **Carolles** | 1.01 | 1.2 |
| **CROZ** | **Crozon** | 1.07 | 0.76 |
| ITXA | Itxassou | 0.25 | 0.25 |
| MONT | Plounéour-Ménez | na | na |
| **TREG** | **Trégastel** | 0.7 | 0.5 |

For the PFOA, 216 of the 394 tested strains showed a tolerance coefficient ≥ 1and 39 strains a coefficient ≥ 1.5. Thirteen strains (3.2 % of the tested bacteria) were highly tolerant to the PFOA and as for the MTBE most of them were isolated from the Carolles, Crozon and Trégastel lichen population (Table 2), by reinforcing a possible correlation between the habit of origin and the ability to degrade POP. As for the MTBE, the highest tolerant strains belonged to the *Pseudomonas, Bacillus, Sphingomonas* and *Paenibacillus* genus, suggesting a recurrent ability of these bacterial species to degrade different POP. On the other hand, we also found that some of the most tolerant strains were affiliated to the genus *Methylobacterium, Erythrobacter, Micrococcus* and *Frigoribacterium*.

In our study, POP tolerance was not correlated with antimicrobial activity as described previously [30]. In fact, the most resistant bacterial strains MONT-RG-20F-10-E-7-02, MONT-RG-20F-R14-05, TREG-RG-20F-10-E-5-01 and TREG-RG-20F-10-E-6-01 did not show a strong POP tolerance. Globally, our results corroborate with the evidence that lichen habitats are a source of POP tolerant and antibiotic resistant strains with antimicrobial activities but that those activities are not strictly correlated. On the other hand, the POP tolerance seems to be highly distributed in habitats with a oil spill history (Table 2).

## Discussion

Our novel culturomic approach on lichen-associated bacteria provides a unique picture and database on diversified bacterial communities naturally colonizing *R. geographicum*. Most of the previous culture-dependent studies on lichens only used classic media allowing to isolate a small fraction of lichen-associated microbes [42]. In addition, only two studies explored the diversity of *R. geographicum*. Firstly, Bjelland *et al*. [20], through a 16S rDNA metabarcoding approach investigated the diversity of bacterial communities associated with *R. geographicum* without combining culturing methods. Secondly, a recent study from Miral *et al*. [63] isolated 24 bacteria strains and 68 fungal isolates from *R. geographicum* by using classical microbiological media. Miral and co-authors did not use antifungal compounds for bacterial isolation and by consequence they isolated a low bacterial diversity. For instance, most of the isolated bacteria belonged to the *Paenibacillus etheri* species. Our study is then the first deep ecological investigation of bacterial communities associated to *R. geographicum*, a lichen undoubtedly difficult to sample from rock surfaces but adapted to extreme environmental conditions. To our knowledge only few works used original media enriched with lichen extract in order to isolate microbes from other lichen species [27, 64]. The “Microbial Dark Matter” [65] might be recovered in laboratory conditions through a reasonable imitation of the habitat where these microbes evolve during their ecological time. In this concern, media developed in our study to mimic *R. geographicum* environment, allowed to isolate bacterial species not identified before on conventional media. Our results claim that specific-host nutrients play a substantial role in isolating rare species. Developing culturing methods on these nutrients is the missing key to build diversified microbial collections from lichens with an undiscussed importance for *eco-evo* studies on holobionts such as lichens. Indeed, among the 699 single clusters amplified and characterized with the 16S marker gene, 19.9 % of bacteria were identified as uncultured, including an unclassified strain and three strains with any matches when blasted on GenBank. Among the 11 whole-genome sequenced strains, 9 were putative novel species. Moreover, the long incubation time allowed to recover strains belonging to the Frankiales order [66] hardly isolated from lichens before. The only study on the *R. geographicum* microbial diversity, Bjelland et al. in 2011 [20], found that bacterial community of *R. geographicum* consisted mainly of Acidobacteria, Proteobacteria (Alpha- and Beta-proteobacteria) and Chloroflexi classes with a poor discrimination at the order level due to a lack of metabarcoding resolution. By culturomics, at the bacterial class level we also found that Alpha-proteobacteria and Beta-proteobacteria are predominant bacterial classes of the *R. geographicum* microbiota but we also isolated a large amount of Actinomycota. Furthermore, in our study we deeply characterized the isolated bacteria at the family and species order and we succeeded to isolate rare species previously known to be adapted to extreme environments (Table 1).

The principal aim of our study was not only to initiate a *R. geographicum*-associated bacterial collection but to utilize this collection to understand the ecological features of bacterial communities in relationship to their habitat of origin. As some of the strains were isolated from media containing antibiotics, we investigated their resistance degree and demonstrated that part of the bacteria-associated with lichens harbor resistance to a large class of antibiotics. Reconstruction of KEGG metabolic pathways confirmed the presence of potential functions protecting bacteria against toxic compounds. For example, we found metabolic pathways related to arsenic reduction in all whole-genome sequenced bacteria (Supplementary Data Set 4). Arsenic can be present in the environments where lichens were recovered; therefore, bacteria might metabolize or transform arsenic from the habitat as described in previous species like *Bacillus* sp. [67]. Interestingly, the most antibiotic resistant bacterial strains (CROZ-RG-20F-R04-06, CROZ-RG-20F-R04-15, MONT-RG-20F-10-E-7-02, MONT-RG-20F-R14-05, TREG-RG-20F-10-E-5-01 and TREG-RG-20F-10-E-6-01), carry sulfur assimilation and oxidation pathways. It has been demonstrated that inhibitors of sulfur assimilation pathways in human bacterial pathogens can act as enhancer of antibiotic therapy by suggesting a close link between the sulfur metabolization and the resistance to antibiotics in bacteria [68]. Together with pathways for arsenic metabolization, sulfur assimilation could also confer to lichen-associated bacteria the ability to resist to antibiotics. On the other hand, antibiotic resistance activities can emerge under extreme environmental conditions and be related to more large resistance mechanisms (i.e. resistance to pollutants or competition among microbial communities in a nutrient poor environment).

AMR gene analyses on the 11 genomes showed resistance mechanisms for antibiotics, such as classes of β-lactamases and resistance-nodulation-division (RND) efflux pumps for strains identified as potential novel species. The antibiotic resistance pathways found in these strains are common in Gram negative bacteria [69] and well-studied in many *Pseudomonas* species [70, 71]. For instance, Multidrug Resistant (MDR) pumps play an important role during the first step of plant colonization in bacterial phytopathogens [72–74]. The relevance of efflux pumps in plant-bacterial interactions was also described in symbiotic bacteria. For example, mutants of *Rhizobium etli*, a mutualistic symbiont of the *Phaseolus vulgaris* bean, with a defective RmrAB efflux pump formed on average 40 % less nodules than the wild-type strain [75]. Taken together, results on different habitats/hosts indicate that antibiotic resistance is a mechanism not exclusively related to protect the bacterium against antimicrobial compounds but also to help the bacterium to adapt and evolve to a particular ecological context. In light of this, the lichenized fungus produces specialized metabolites with antimicrobial properties (e.g. depsidones, depsides and dibenzofurans) [76]; these compounds are a serious source of abiotic stress for the bacterial communities associated to the lichens. *R. geographicum* produced a vulpinic acid derivative, rhizocarpic acid, which has shown moderate antimicrobial activity (MIC range of 32-64 µg mL^-1^) against various multi-resistant *Staphyloccocus aureus* strains [77]. The presence of RND efflux pumps and other antimicrobial resistance pathways can then confer to the bacterial microbiota the arsenal to adapt to the lichen [37]. Furthermore, the RND efflux pumps are involved in Polyaromatic Hydrocarbons (PAHs) degradation [78, 79]; a group of organic compounds which are highly toxic and contaminate terrestrial and aquatic environments. RND-type efflux pump (EmhABC) has been described previously [78, 80] as extruding hydrophobic antibiotics and PAHs, including phenanthrene, anthracene, and fluoranthene [78, 81]. Taking into consideration that in our study *R. geographicum* was sampled in littorals with a history of oil spill, the resistome (the complex of AMR genes) might be emerged in lichen-associated bacteria in response to pollutants to help bacteria resilience in contaminated habitats [82, 83]. In accordance to our hypothesis, several studies revealed that the genus *Pseudomonas* was dominant in PAH-contaminated sites and that hydrocarbon degradation ability and antibiotic resistance were strongly correlated in these bacteria [84]. In addition, the emergence of AMR genes was demonstrated to be 15 times higher in contaminated sites [85] [86].

Our results demonstrated that on average 60 %, for MTBE, and 54 % for PFOA, of the strains tested for POP resistance were able to grow and tolerate POP (tolerance coefficient ≥ 1). But unlike what was described before, resistance to POP was not correlated to antibiotic resistance, when considering the most tolerant strains ((tolerance coefficient ≥ 1.5 and 2). Interestingly, strains with the highest tolerant coefficient were isolated from lichens that experienced a history of oil spill. This observation indicates that pollution and environmental constraints can select from bacterial lineages with the ability to tolerant and probably metabolize pollutants. Therefore, mass isolation and characterization of strains from these particular habitats can help in identifying novel microbial candidates for biotechnological purposes. In light of this, culturomics needs to be developed for selecting highly pollutants degrading strains. It is worth to notice, that some POP tolerant strains were not recovered from habitats with oil spill history. In this concern, as lichens are known to be extreme habitats for microbes, resistance and tolerance mechanisms might evolve without a precise pollutant and the diversity of the strains and their different resistances widen the scope of biotic stresses that the holobiont can handle.

As PAHs and the POP tested in our study are organic substances characterized by their persistence, toxicity, mobility on long distances and their bio-accumulative nature [87], the bacterial collection we established is relevant under a biotechnological level. In fact, nowadays, these compounds represent a genuine threat to wild life preservation and to human health with an urgent need to counterbalance this pollution. Bioremediation has generally been considered a sustainable approach to managing petroleum hydrocarbon contaminated soils. There has been an increasing focus on ‘green’ and ‘sustainable’ remediation and an international standard ISO18504:2017 Soil quality – Sustainable remediation was recently published (2017) [82]. Based on the POP tolerance of the bacterial strains we isolated, this is a first step to develop Synthetic Microbial Communities very effective in POP degradation and then a unique tool to manage the bioremediation of polluted sites.

## Supporting information

Data Set

## Acknowledgements

This study was funded by Défi Scientifique 2022 of the University of Rennes 1 and the INRAE-SPE ECOSYMB project. We particularly thank Bruno Marquer technician at the IGEPP laboratory (INRAE, Rennes) for his technical support during the antibiotic resistance assay and POP tolerance assay. We also thank Julien Fournet for his kind help in the molecular biology strain characterization. Finally, we thank Léa Frachon from University of Zurich, for her advices in manuscript and images adjustments.

## Supplementary Material

**Supplementary Information** including supplementary Methods, Tables and Figures.

**Supplementary Data Set 1**. Bacterial Clusters sharing 100% of sequence homology with two or more strains after CD-HIT analysis.

**Supplementary Data Set 2**. Taxonomy of the 696 bacterial unique clusters identified after CD-HIT trimming.

**Supplementary Data Set 3**. 16S sequences of strains isolated on media containing antibiotics.

**Supplementary Data Set 4**. Heatmap on the KEGG metabolic pathways found on the strains that were whole-genome sequenced. Antibiotic resistant strains are marked with “*”. Gradient color from with to red reflects the completeness of the metabolic pathway.

**Supplementary Data Set 5**. Protein classes found by using the AMRFinder Plus pipeline to be associated with antibiotic resistance or resistance to heavy metals. The analysis was performed on the 9 genomes of strains described to carry a broad spectrum of resistance to antibiotics. Protein sequences are reported.

**Supplementary Data Set 6**. LSmeans estimated on the POP tolerance of each strain.

## Supplementary Information for

### SUPPLEMENTARY METHODS

#### Media used for bacteria isolation from *R. geographicum*

Media codes are indicated into parentheses after the medium name. <u>This code has been added to the bacterial strains and it represents a code belonging to a larger list of media developed at the IGEPP Laboratory (INRAE, Bretagne, France)</u>.

Product references are reported to allow the repeatability of the preparation.

#### Nutrient Agar (01B)

##### Per Liter of medium

**Per Liter of medium:**
|  |  |
| --- | --- |
| Osmotic water | 1 L |
| Beef extract (VWR, Reference: 1.03979.0500 500 g) | 3.0 g |
| Peptone (GICO, Reference: 211677 500 g) | 5.0 g |
| Glucose (Sigma, Reference: 16301 1 kg) | 1.0 g |
| Yeast extract (BD, Reference 212750 500 g) | 0.5 g |
| Bacto Agar (BD, Reference 214010 500 g) | 15 g |

After autoclaving, add 100 mg.L-1 of cycloheximide (Sigma, Reference: 239764).

#### Tryptic Soy Agar Minimal (08B)

##### Per Liter of medium

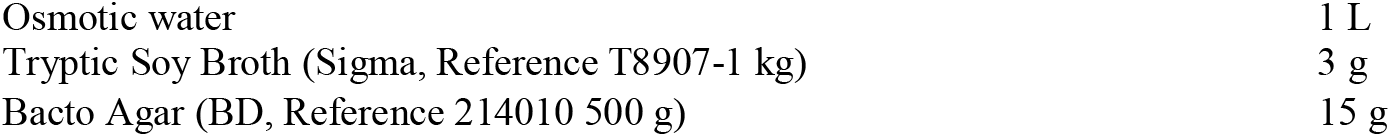

After autoclaving, add 100 mg.L-1 of cycloheximide (Sigma, Reference: 239764).

#### Macroalgae-based medium (14B)

##### Per Liter of medium

**Per Liter of medium:**
|  |  |
| --- | --- |
| Osmotic water | 900 mL |
| K<sub>2</sub>HPO<sub>4</sub>, (Sigma, Reference: P3786 500 g) | 3.5 g |
| KH<sub>2</sub>PO<sub>4</sub>, (VWR, Reference: 1 kg) | 1.0 g |
| MgSO<sub>4</sub> *7H<sub>2</sub>O, (Sigma, Reference: 230391 1 kg) | 1.5 g |
| Glucose (Sigma, Reference: 16301 1 kg) | 0.5 g |
| Bacto Agar (BD, Reference 214010 500 g) | 15 g |

check pH (∼6)

After autoclaving, add 50 mL of each macroalgae filtrate (50 ml of the *Ascophyllum nodosum* filtrate and 50 mL osf the *Laminaria digitata* filtrate) described in the Material and Methods section of the main text. Also add 100 mg.L-1 of cycloheximide (Sigma, Reference: 239764).

#### Lichen algal-based medium (15B)

##### Per Liter of medium

**Per Liter of medium:**
|  |  |
| --- | --- |
| Osmotic water | 900 mL |
| K<sub>2</sub>HPO<sub>4</sub>, (Sigma, Reference: P3786 500 g) | 3.5 g |
| KH<sub>2</sub>PO<sub>4</sub>, (VWR, Reference: 1 kg) | 1.0 g |
| MgSO<sub>4</sub> *7H<sub>2</sub>O, (VWR, Reference: 1 kg) | 1.5 g |
| Glucose Sigma, Reference: 16301 1 g) | 0.5 g |
| Bacto Agar (BD, Reference 214010 500 g) | 15 g |

check pH (∼6)

After autoclaving, add 50 mL of each macroalgae filtrate (50 ml of the *Ascophyllum nodosum* filtrate and 50 mL of the *Laminaria digitata* filtrate) described in the Material and Methods section of the main text. In addition, add 5 ml of lichen filtrate prepared as described in the main text. Also add 100 mg.L-1 of cycloheximide (Sigma, Reference: 239764).

#### Lichen algal-based minimal medium (16B)

##### Per Liter of medium

**Per Liter of medium:**
|  |  |
| --- | --- |
| Osmotic water | 800 mL |
| Sea water | 95 mL |
| Bacto Agar (BD, Reference 214010 500 g) | 15 g |

check pH (∼6)

After autoclaving, add 50 mL of each macroalgae filtrate (50 ml of the *Ascophyllum nodosum* filtrate and 50 mL of the *Laminaria digitata* filtrate) described in the Material and Methods section of the main text. In addition, add 5 mL of lichen filtrate prepared as described in the main text. Also add 100 mg.L-1 of ycloheximide (Sigma, Reference: 239764).

#### Lichen microalgal-based minimal medium (17B)

##### Per Liter of medium

**Per Liter of medium:**
|  |  |
| --- | --- |
| Osmotic water | 850 mL |
| Sea water | 95 mL |
| Bacto Agar (BD, Reference 214010 500 g) | 15 g |

check pH (∼6)

After autoclaving, add 50 mL of the *Coccomyxa viridis* filtrate obtained as described in the material and methods section of the main text. In addition, add 5ml of lichen filtrate prepared as described in the main text. Also add 100 mg.L-1 of cycloheximide (Sigma, Reference: 239764).

#### Algae antibiotic-based medium (18F)

##### Per Liter of medium

**Per Liter of medium:**
|  |  |
| --- | --- |
| Osmotic water | 800 mL |
| Papaic Digest of Soybean Meal (Gibco, Reference 2436 500 g) | 2.0 g |
| Dextrose (Glucose C<sub>6</sub>H<sub>12</sub>O<sub>6</sub>) (Sigma, Reference 16301 1 kg) | 1.0 g |
| Sea water | 100 ml |
| Bacto Agar (BD, Reference 214010 500 g) | 15 g |

check pH (∼6)

After autoclaving, add 50 mL of each macroalgae filtrate (50 mL of the *Ascophyllum nodosum* filtrate and 50 mL of the *Laminaria digitata* filtrate) described in the Material and Methods section of the main text. Also, add 100 mg.L-1 of chloramphenicol (Sigma, Reference: C0378-100g), 300 mg.L-1 of streptomycin (Sigma, Reference: S6501) and 100 mg.L-1 penicillin (Sigma, Reference: 1504489).

#### Lichen algal antibiotic-based minimal medium (19F)

##### Per Liter of medium

**Per Liter of medium:**
|  |  |
| --- | --- |
| Osmotic water | 800 mL |
| Sea water | 95 ml |
| Bacto Agar (BD, Reference 214010 500 g) | 15 g |

check pH (∼6)

After autoclaving, add 50 mL of each macroalgae filtrate (50 mL of the *Ascophyllum nodosum* filtrate and 50 mL of the *Laminaria digitata* filtrate) described in the Material and Methods section of the main text. In addition, add 5 mL of lichen filtrate prepared as described in the main text. Also, add 100 mg.L-1 of chloramphenicol (Sigma, Reference: C0378-100g), 300 mg.L-1 of streptomycin (Sigma, Reference: S6501) and 100 mg.L-1 penicillin (Sigma, Reference: 1504489).

#### Lichen algal antibiotic-based medium (20F)

|  |  |
| --- | --- |
| Osmotic water | 800 mL |
| Mycological peptone (Sigma 77199-500 g) | 5 g |
| Magnesium sulphate (MgSO<sub>4</sub>) (Sigma, Reference: 77199-500 g) | 2.5 g |

check pH (∼6)

After autoclaving, add 50 mL of each macroalgae filtrate (50 mL of the *Ascophyllum nodosum* filtrate and 50 mL of the *Laminaria digitata* filtrate) described in the Material and Methods section of the main text. In addition, add 5 mL of lichen filtrate prepared as described in the main text. Also, add 100 mg.L-1 of chloramphenicol (Sigma, Reference: C0378-100g), 300 mg.L-1 of streptomycin (Sigma, Reference: S6501) and 100 mg.L-1 penicillin (Sigma, Reference: 1504489).

### Bacterial identification

To characterize the bacterial strain collection, we amplified a portion of the 16S rRNA gene. Universal primers 515F (5′-GTGCCAGCMGCCGCGGTAA-3′) and 806R (5′-GGACTACVSGGGTATCTAAT-3′) [1] were used in 24 µL of PCR mix performed with 5X GoTaq® DNA Polymerase (Promega) kit. 2 µL of the extracted DNAs were added in the mix. PCR conditions are as follows: 95°C for 5 min and 35 cycles at 95°C for 5 s, 55°C for 30 s, 72°C for 60 s with a final extension at 72°C for 5 min. PCR amplicons were verified by loading 5 µL of each product on a 1 % w/v agarose gel. Products showing the suitable band size were then purified by adding 0.5 µL of Antarctic phosphatase, 0.5 µL of Exonuclease I and 1 µL of Phosphatase Buffer to 10 µL of PCR product. Sample mixtures were then incubated for 30 min at 37°C and 10 min at 90°C to deactivate the enzyme. Purified PCR products were sequenced at the Macrogen Europe BC (Amsterdam, Netherlands). Chromatograms were individually analyzed with Chromas Lite v 2.6.6 (Technelysium Ltd.) was used for manual corrections of the obtained raw sequences. and used to build a multi-FASTA file. To remove the clonal sequences and group unique sequence clusters, we run the Cd-hit software [2] with the following parameters -c 1 -aS 1 -d 50000 -T 10. Unique bacterial clusters were then integrated into Blastn with parameters: -evalue 0.000001 -num_threads 10 - max_target_seqs 20 -outfmt. A homemade perl script was then used to reconstruct a taxonomy table represent the phylogenetic affiliation of the strains. Bar plots representing the diversity of the bacterial strains were performed with ggplot2 [3].

#### Whole-genome Sequencing and bioinformatics

Nine bacterial strains showing the highest antibiotic resistance profile (Supplementary Table S3) and two strains whose 16S sequence fragment did not match in GenBank (CARO-RG-8B-R23-01 and CARO-RG-8B-R24-01) were whole genome sequenced. Prior DNA extraction, strains were grown from 2 to 4 days (depending on the growth rate of the strain) in TSB medium. At the stationary phase, bacterial cultures were centrifuged for 10 min at 5000 rpm and the supernatant was discarded. Bacterial pellet was then washed in sterilized MQ water and further centrifuged for 10 min. The collected pellet was then employed for DNA extraction by using the DNeasy Blood & Tissue Kit of Qiagen. A RNase step (with the Invitrogen™ RNase A 20 mg/mL) was added to the kit protocol to obtain RNA-free DNAs. For this 5 µL of RNaseA were added into the samples that were incubated at 65°C for 20 min. The concentration and purity of DNAs were estimated with spectomethry. DNA degradation level was checked by running 3 µL of the extracted DNAs on a 1 % w/v agarose gel. Genomic DNA libraries were performed with the Short-Insert Library protocol and sequenced with the DNBseq™ platform (PE150 for sequencing length) at the BGI sequencing facility (Shenzhen, China).

Raw reads in the Fastq file format were filtered by removing adaptor sequenced, contaminations and low-quality reads by BGI. Clean reads were then annotated and assembled by using the Bacannot framework https://bacannot.readthedocs.io/en/latest/#index--page-root. KEGG KO annotation was performed with KofamScan version 1.3.0 [4]. Taxonomy affiliation on the whole genome sequences was performed with TYGS (Type Strain Genome Server) https://tygs.dsmz.de/ [5].

Metabolic functional analysis on the nine bacterial genomes was performed. First, for each bacterial genome, a list of biochemical reactions was inferred from its proteic sequences with Carveme software [6]. Then, each reaction was classified into metabolic pathways by querying the BIGG database [7]. To compare the presence of metabolic pathways and reactions, presence-absence matrices were computed thanks to ad-hoc programs developed with the Met4j library ^1^.

## SUPPLEMENTARY TABLES

**Supplementary Table S1.** Names and GPS coordinates (expressed in degrees) of the 24 sites on which *Rhizocarpon geographicum*.

| **Population name** | **Locality** | **Altitude (m)** | **Latitude** | **Longitude** |
| --- | --- | --- | --- | --- |
| CROZ-RG-R01 | Crozon | 30 | 48.2333333 | -4.565277777777 |
| CROZ-RG-R02 | Crozon | 50 | 48.2330556 | -4.56638888888888 |
| CROZ-RG-R03 | Crozon | 40 | 48.2319444 | -4.57027777777777 |
| CROZ-RG-R04 | Crozon | 10 | 48.2336111 | -4.56555555555555 |
| BAUL-RG-R05 | Baulon | 60 | 47.9713889 | -1.8975 |
| BAUL-RG-R06 | Baulon | 60 | 47.9713889 | -1.8975 |
| BAUL-RG-R07 | Baulon | 80 | 47.9722222 | -1.89749972222222 |
| BAUL-RG-R08 | Baulon | 80 | 47.9725 | -1.89749972222222 |
| ITXA-RG-R09 | Itxassou | 610 | 43.3052778 | -1.42888888888888 |
| ITXA-RG-R10 | Itxassou | 610 | 43.3025 | -1.42805555555555 |
| ITXA-RG-R11 | Itxassou | 620 | 43.3022222 | -1.42805555555555 |
| ITXA-RG-R12 | Itxassou | 660 | 43.3016667 | -1.4325 |
| MONT-RG-R13 | Plounéour-Ménez | 390 | 48.4055556 | -3.90999999999999 |
| MONT-RG-R14 | Plounéour-Ménez | 370 | 48.8369444 | -3.50388888888888 |
| MONT-RG-R15 | Plounéour-Ménez | 380 | 48.4058333 | -3.90944444444444 |
| MONT-RG-R16 | Plounéour-Ménez | 380 | 48.4055556 | -3.90916666666666 |
| TREG-RG-R17 | Trégastel | 10 | 48.8366667 | -3.50388888888888 |
| TREG-RG-R18 | Trégastel | 20 | 48.8369444 | -3.50388888888888 |
| TREG-RG-R19 | Trégastel | 20 | 48.8372222 | -3.5 |
| TREG-RG-R20 | Trégastel | 10 | 48.8372222 | -3.50194444444444 |
| CARO-RG-R21 | Carolles | 70 | 48.7441667 | -1.56944444444444 |
| CARO-RG-R22 | Carolles | 60 | 48.7444444 | -1.56972222222222 |
| CARO-RG-R23 | Carolles | 60 | 48.7463889 | -1.57055555555555 |
| CARO-RG-R24 | Carolles | 60 | 48.7461111 | -1.57055527777777 |

**Supplementary Table S2.** Mass of *R. geographicum* lichen used for bacterial isolation.

| **Population name** | **Mass (mg)** |
| --- | --- |
| CROZ-RG-R01 | 80.2 |
| CROZ-RG-R02 | 250 |
| CROZ-RG-R03 | 72.9 |
| CROZ-RG-R04 | 53.5 |
| BAUL-RG-R05 | 33 |
| BAUL-RG-R06 | 80 |
| BAUL-RG-R07 | 33.9 |
| BAUL-RG-R08 | 40.1 |
| ITXA-RG-R09 | 28.9 |
| ITXA-RG-R10 | 37.4 |
| ITXA-RG-R11 | 43.9 |
| ITXA-RG-R12 | 27.4 |
| MONT-RG-R13 | 30.2 |
| MONT-RG-R14 | 32.8 |
| MONT-RG-R15 | 37.6 |
| MONT-RG-R16 | 16.5 |
| TREG-RG-R17 | 164.7 |
| TREG-RG-R18 | 250.7 |
| TREG-RG-R19 | 525.4 |
| TREG-RG-R20 | 449.6 |
| CARO-RG-R21 | 42.2 |
| CARO-RG-R22 | 36.2 |
| CARO-RG-R23 | 48.4 |
| CARO-RG-R24 | 45.4 |

**Supplementary Table S3.**
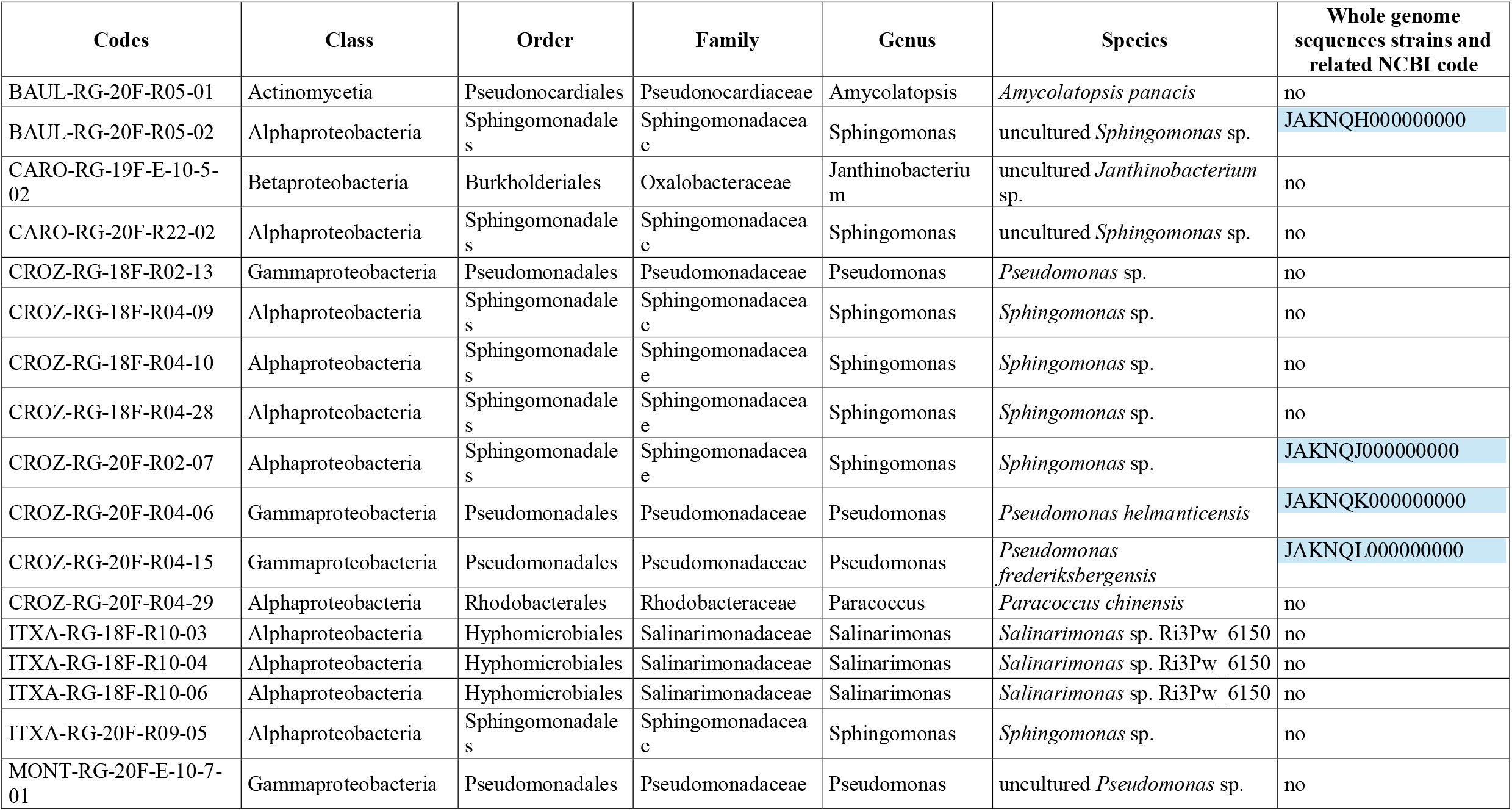
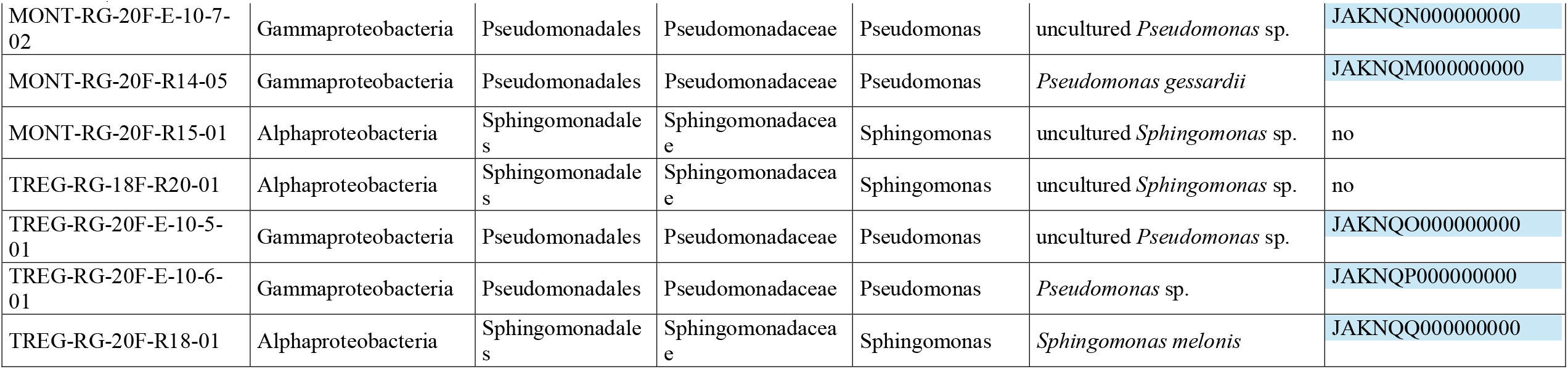
Bacteria isolated on media containing antibiotics and used for the antibiotic susceptibility test. Taxonomic affiliation was assigned by blastn.

**Supplementary Table S4.** Antibiotics and relative concentrations (µg/mL) used for the antibiotic susceptibility test.

| Antibiotics | Concentration ($\mu\text{g/mL}$) |
| --- | --- |
| Gentamicin | 5 |
| Ampicillin | 100 |
| Streptomycin | 150 |
| Kanamycin | 50 |
| Carbenicillin | 100 |
| Penicillin G | 100 |
| Tetracycline | 12 |
| Rifampicin | 50 |
| Vancomycin | 10 |
| Chloramphenicol | 100 |
| Colistin | 5 |
| Cefalexin | 50 |

**Supplementary Table S5.** Statistics of clean genomic data obtained after fastq trimming

| Sample name | Clean Reads | Clean Base | Read Length | Q20 (%) | GC (%) |
| --- | --- | --- | --- | --- | --- |
| BAUL-RG-20F-R05-02 | 4,203,811 | 1,261,143,300 | PE150 | 93.85 | 63.69 |
| CARO-RG-8B-R23-01 | 4,057,029 | 1,217,108,700 | PE150 | 93.51 | 69.56 |
| CARO-RG-8B-R24-01 | 4,052,146 | 1,215,643,800 | PE150 | 94.17 | 64.73 |
| CROZ-RG-20F-R02-07 | 4,205,378 | 1,261,613,400 | PE150 | 93.36 | 67.13 |
| CROZ-RG-20F-R04-06 | 4,126,825 | 1,238,047,500 | PE150 | 93.94 | 58.73 |
| CROZ-RG-20F-R04-15 | 4,131,243 | 1,239,372,900 | PE150 | 94.31 | 58.59 |
| MONT-RG-20F-10-7-02 | 4,028,310 | 1,208,493,000 | PE150 | 94.52 | 59.94 |
| MONT-RG-20F-R14-05 | 4,035,850 | 1,210,755,000 | PE150 | 93.9 | 59.94 |
| TREG-RG-20F-10-5-01 | 4,020,368 | 1,206,110,400 | PE150 | 93.89 | 59.88 |
| TREG-RG-20F-10-6-01 | 4,014,265 | 1,204,279,500 | PE150 | 93.98 | 59.85 |
| TREG-RG-20F-R18-01 | 4,189,150 | 1,256,745,000 | PE150 | 93.4 | 65.03 |

**Supplementary Table S6.** Analysis of Deviance Table (Type II Wald chisquare tests) on the antibiotic resistance coefficient of the 24 strains selected for antibiotic resistance assays. Phase indicates the exponential or stationary phase at which strains have been tested. The phase and the strain *factor* were analyzed as fixed variables. *P* value adjustment was estimated with Tukey method for comparing a family of 48 estimates.

|  | *Chisq* | *Df* | *Pr(>Chisq)* |
| --- | --- | --- | --- |
| **Gentamicin** |  |  |  |
| Strain | 393.653 | 23 | **<2.2e-16** |
| Phase | 28.092 | 1 | **1.157e-07** |
| Strain:Phase | 110.249 | 23 | **2.280e-13** |
| **Ampicillin** |  |  |  |
| Strain | 171.0278 | 23 | **<2.2e-16** |
| Phase | 5.5523 | 1 | 0.01846 |
| Strain:Phase | 80.3465 | 23 | **2.79e-08** |
| **Streptomycin** |  |  |  |
| Strain | 181.1994 | 23 | **<2.2e-16** |
| Phase | 0.9611 | 1 | 0.3269 |
| Strain:Phase | 103.8320 | 23 | **3.047e-12** |
| **Kanamycin** |  |  |  |
| Strain | 195.614 | 23 | **<2.2e-16** |
| Phase | 29.224 | 1 | **6.446e-08** |
| Strain:Phase | 109.919 | 23 | **2.607e-13** |
| **Carbenicillin** |  |  |  |
| Strain | 185.5520 | 23 | **<2.2e-16** |
| Phase | 0.0289 | 1 | 0.8651 |
| Strain:Phase | 76.0049 | 23 | **1.391e-07** |
| **Penicillin** |  |  |  |
| Strain | 173.1999 | 23 | **<2.2e-16** |
| Phase | 2.1399 | 1 | 0.14351 |
| Strain:Phase | 34.4238 | 23 | 0.05927 |
| **Tetracycline** |  |  |  |
| Strain | 458.8361 | 23 | **<2.2e-16** |
| Phase | 1.6341 | 1 | 0.2011 |
| Strain:Phase | 77.5157 | 23 | **7.979e-08** |
| **Rifampicin** |  |  |  |
| Strain | 520.8060 | 23 | **<2.2e-16** |
| Phase | 0.7714 | 1 | 0.3798 |
| Strain:Phase | 78.6529 | 23 | **5.239e-08** |
| **Vancomycin** |  |  |  |
| Strain | 568.624 | 23 | **<2.2e-16** |
| Phase | 11.026 | 1 | **0.0008983** |
| Strain:Phase | 121.100 | 23 | **2.638e-15** |
| **Chloramphenicol** |  |  |  |
| Strain | 519.4214 | 23 | **<2.2e-16** |
| Phase | 5.5191 | 1 | 0.01881 |
| Strain:Phase | 175.8313 | 23 | **2,00E-16** |
| **Colistin** |  |  |  |
| Strain | 507.401 | 23 | **<2.2e-16** |
| Phase | 24.529 | 1 | **7.321e-07** |
| Strain:Phase | 96.928 | 23 | **4.750e-11** |
| **Cefalexin** |  |  |  |
| Strain | 162.230 | 23 | **<2.2e-16** |
| Phase | 18.782 | 1 | **1.466e-05** |
| Strain:Phase | 103.098 | 23 | **4.090e-12** |

## SUPPLEMENTARY FIGURES

**Figure S1.**
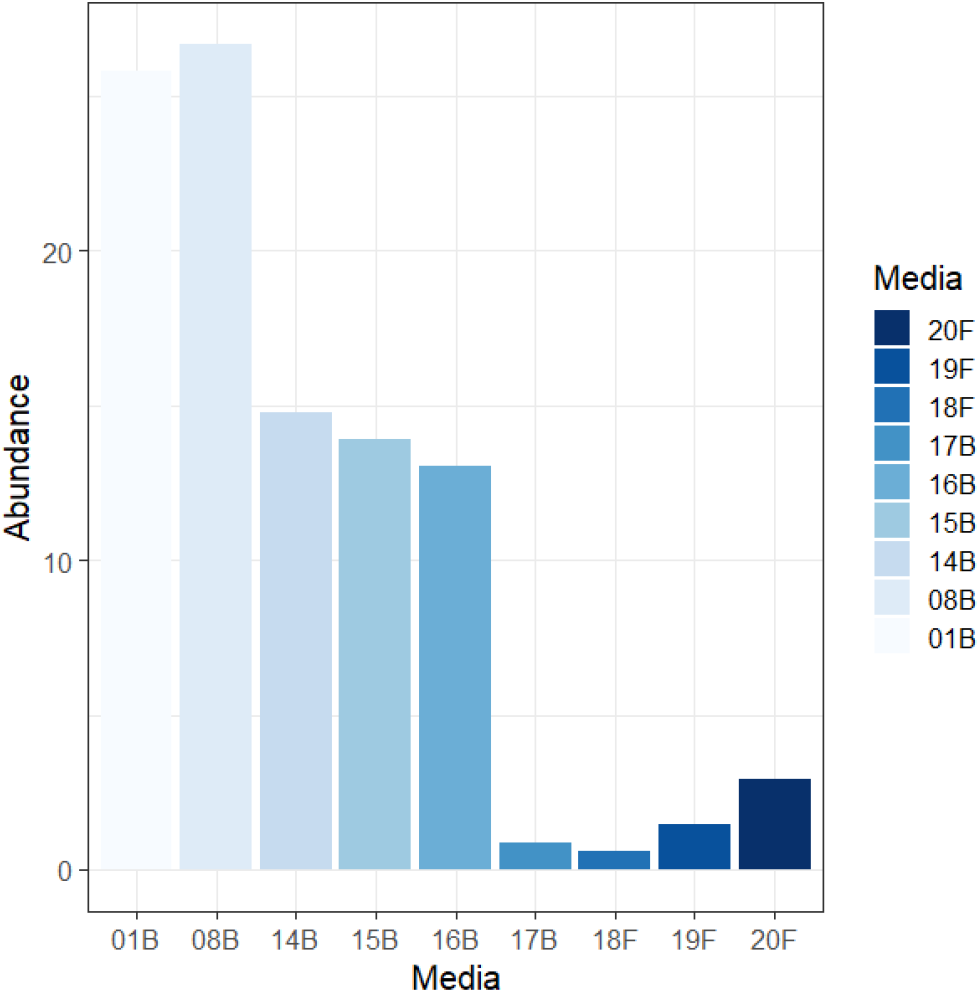
Number of species isolated for each medium. Bar plot is based on the abundance of each bacterial species (expressed in %) calculated by dividing the number of species recovered on each medium by the number of the total species recovered. Bleu scale color represents the different media used for bacterial isolation (see Supplementary Methods). Bar plot clearly shows that as observed for the abundance of the bacterial isolated for each medium, the 01B and 08B media recovered the maximum of the bacterial diversity.

**Figure S2.**
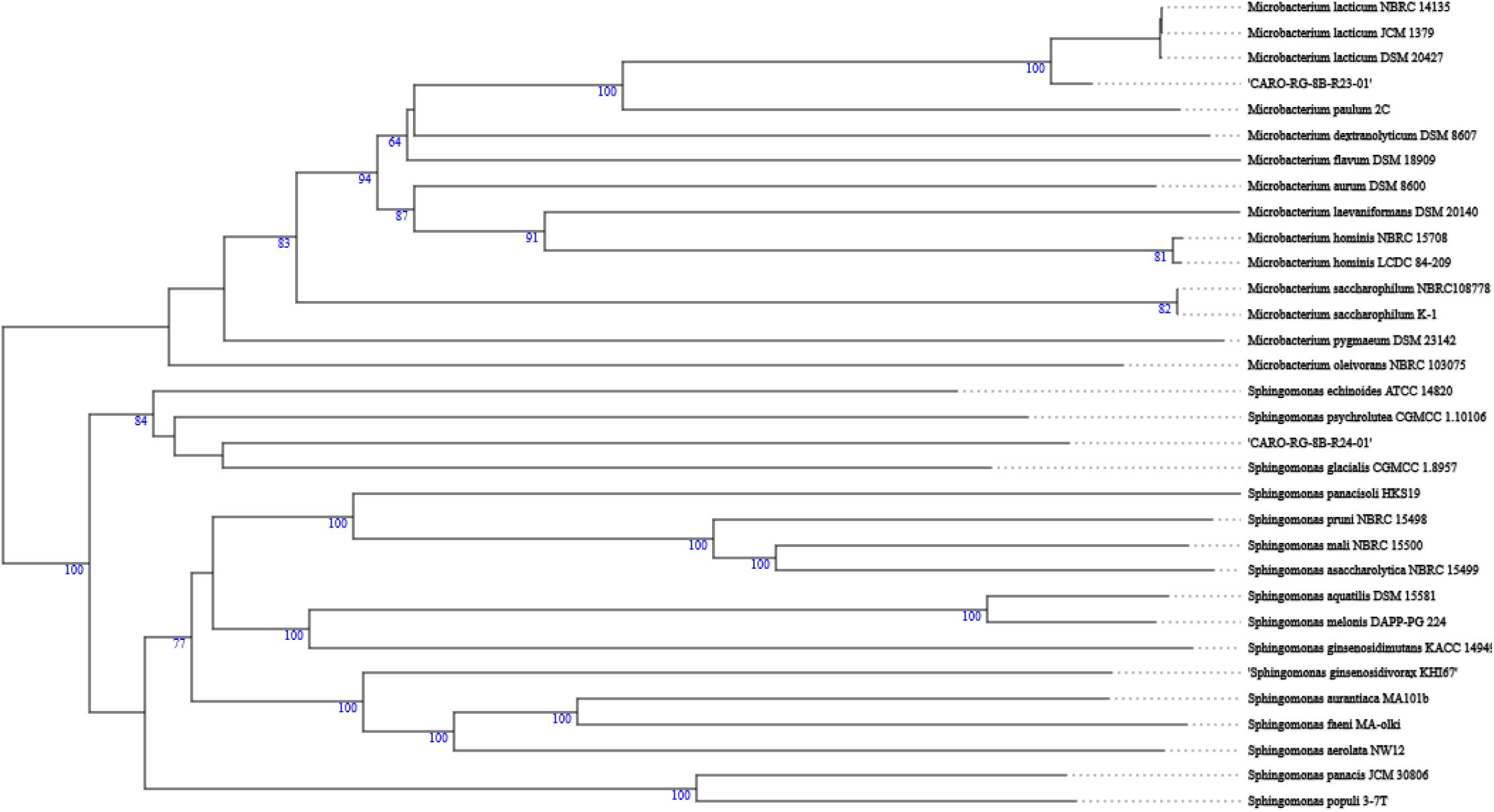
Phylogenetic tree based on the whole genome of the CARO-RG-8B-R24-01 and CARO-RG-8B-R23-01 strains obtained by the Type Strain Genome Service (TYGS).

**Figure S3.**
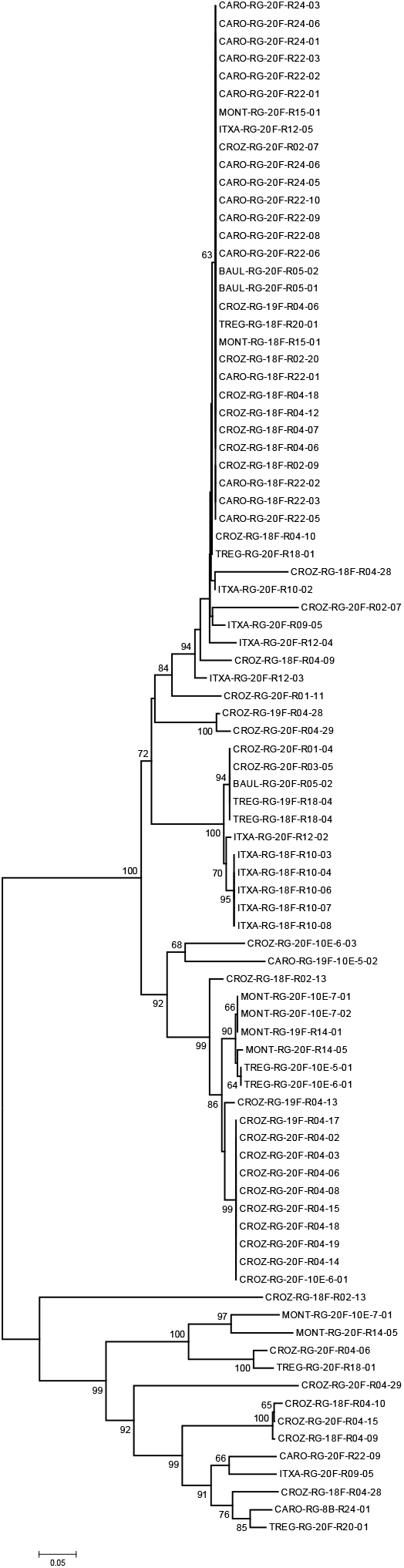
Phylogenetic tree based on the 16S sequences of strains isolated from media containing antibiotics.

**Figure S4.**
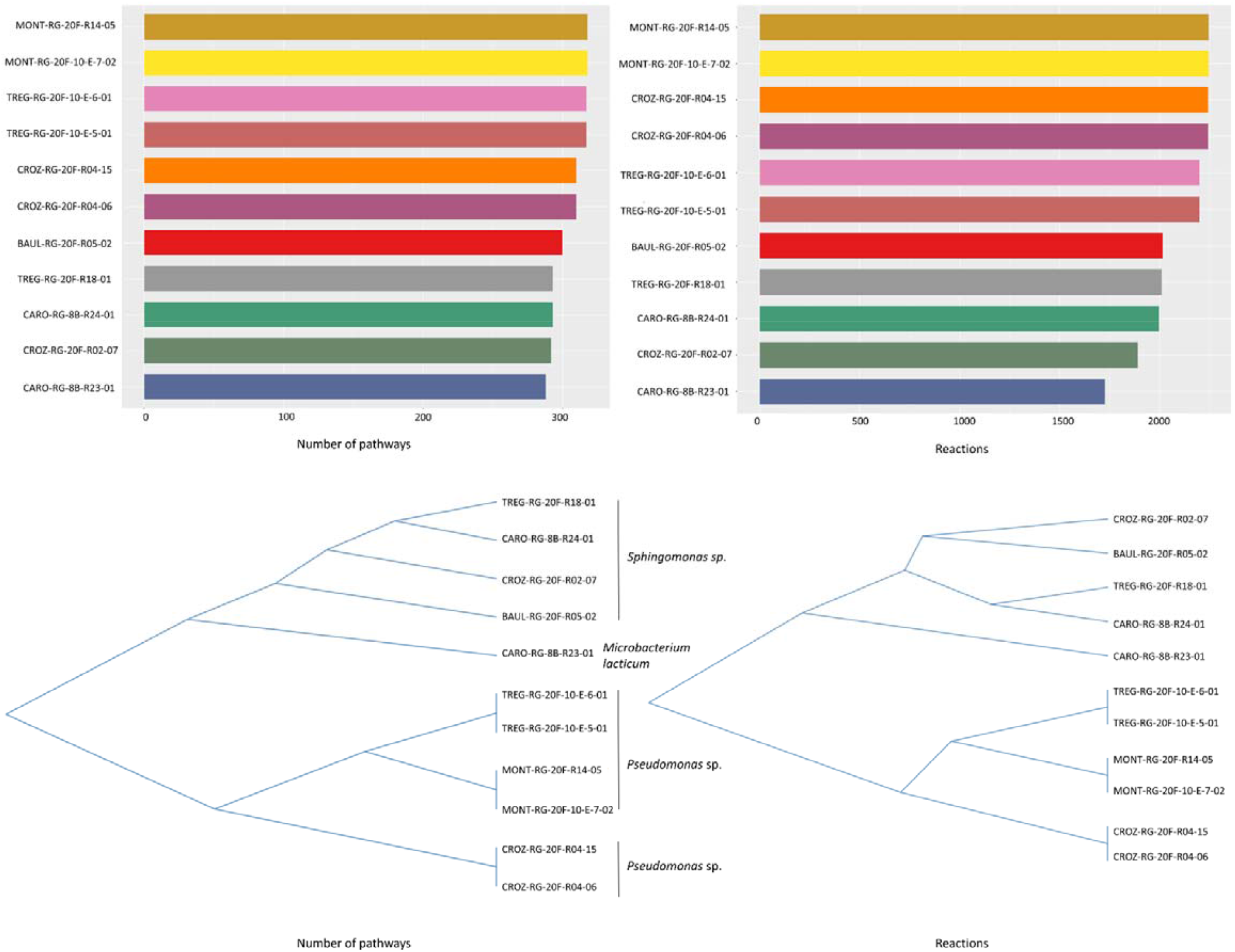
Network analysis based on genome functional annotation. Number of pathways (top left barplot) and number of reactions (top right barplot) based on the annotations of the 11 genomes. Clustering analyses based on the presence/absence of biological pathways (bottom left) and biological reactions (bottom right). The functional clustering is coherent with the taxonomical annotation of the strains. Plots were performed thanks to the MeCompR shiny application^2^.

**Figure S5.**
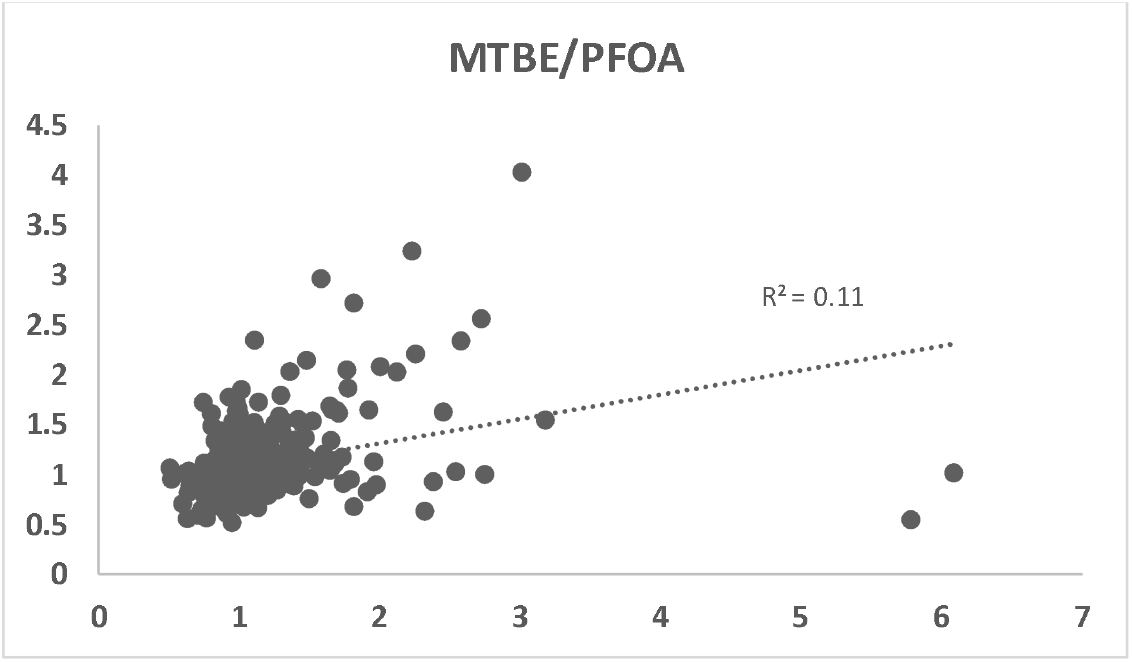
Correlation between Lsmeans values obtained from the POP tolerance experiment. *x*-axis represents Lsmeans values obtained from strains grew in presence of PFOA and *y*-axis represents Lsmeans values obtained from strains grew in presence of MTBE. The *R*^*2*^ representing the correlation between the two variables is also reported.

## Footnotes

1 https://forgemia.inra.fr/metexplore/met4j

2 https://lipm-gitlab.toulouse.inra.fr/lcottret/mecompr

## Notes

### Competing Interest Statement

The authors have declared no competing interest.

